# Unique evolutionary radiation of odorant receptors in birds

**DOI:** 10.1101/2025.11.14.688425

**Authors:** Robert J. Driver, Mona A. Marie, Victoria J. Ko, Priyanka Meesa, Wanting Sun, Renee J. Li, Michael S. Brewer, Hsiu-Yi Lu, Kevin F. P. Bennett, Marco Sollitto, Giulio Formenti, Ichie Ojiro, Nivritti E. Mantha, Hiroaki Matsunami, Christopher N. Balakrishnan

## Abstract

Odorant receptors (ORs) form among the largest gene families in vertebrates; most mammals have hundreds of intact OR genes. Although birds display diverse behavior and ecology, they were long thought to rely minimally on olfaction. We reexamine this idea by using genomic data to uncover diverse and variable OR repertoires in birds, including the nocturnal kiwi (*Apteryx mantelli*), which possesses the largest number of ORs known (3,750). We reveal that widespread gene conversion is shaping the evolution of bird ORs, reflecting a novel mechanism for generating functional diversity. We show that ORs are expressed in avian olfactory sensory neurons and show responses to defined, ecologically relevant odorants. Our results demonstrate the importance and functional mechanisms underlying olfaction in birds.

---

In 1834, Charles Darwin tested if birds could smell by hiding a piece of meat in a folded white paper. When Darwin showed the folded paper to vultures, they showed no interest in the concealed meat. However, when the paper was torn open and the food was revealed, the vultures began to flap and peck excitedly. Darwin wrote that “it would have been quite impossible to have deceive[d] a dog [in this way] (*1*).” This observation aligned with other early studies concluding that olfactory signals were of limited use to birds (*1–3*). In modern times, behavioral work in birds has demonstrated promising roles for olfaction in foraging, locating nest sites, seed caching behavior, and species recognition, among other behaviors (*4–7*). Additionally, specific bird species rely on olfaction for foraging, including the turkey vulture (*Cathartes aura*) and many seabirds (order Procellariiformes) (*8*, *9*). However, how the bird olfactory system functions at the molecular or cellular level remains largely unexplored.

In vertebrates, odors are primarily detected with odorant receptors (ORs), a gene family of G protein-coupled receptors expressed in the olfactory sensory neurons (OSNs) of the olfactory epithelium (*10*, *11*). To accommodate the incredible variety of odorants in nature, ORs constitute one of the largest gene families in vertebrates, for example, there are more than 2,000 OR genes in elephants (*12*, *13*). Early studies using Southern blot and PCR suggested that some birds may possess hundreds of ORs (*14*). We found that genomic OR repertoires in long read assemblies of three bird species ranged from 50 to over 350 intact ORs per species, substantially increasing the number of known ORs in birds (*15*). Therefore long read assemblies provide a unique opportunity to accurately survey ORs across the avian tree of life.

Despite the potential importance of olfaction in bird behavior and ecology, there is little understanding of the evolutionary diversification and expression of ORs, and no information regarding avian OR response properties. Here, we aimed to test whether bird ORs are functional in the avian olfactory system and to reevaluate the historical view that olfaction is largely irrelevant to bird life history. First, using long read, publicly available assemblies (a majority from the Vertebrate Genomes Project Phase I, *16*) we document patterns of OR diversity across the avian tree of life and the evolutionary processes underlying this diversity. We then localized OSNs in chicken (*Gallus gallus*) olfactory epithelium, and analyzed the extent to which individual OR expression was localized to the OSNs. Finally, we determined that bird ORs and OSNs are capable of responding to odor, and identified specific odorants that trigger responses.

## Results

### Characterization of bird odorant receptor repertoires

To understand OR repertoire size and diversity across bird species, we used a BLAST search with known OR sequences as a query to detect complete, putatively functional OR genes in 148 bird genome assemblies from across the phylogeny (see Methods for details, Fig. 1, A and B). We previously demonstrated that short-read genome assemblies consistently underestimate OR gene counts due to high sequence similarity between many bird ORs (fig. S1, *15*). To more consistently capture OR genes, we selected only genome assemblies using long read sequencing technology, with a minimum contigN50 size of 7 Mb. Of these long read genomes, 76 of 148 (51.35%) were from Vertebrate Genomes Project Phase I (*16*). We found a wide range of OR repertoires among birds, in many cases surpassing humans and other mammals (Fig. 1B). We found an average of 251.9 ORs per species, with a median of 146.5 ORs per species. We found an incredible range of OR repertoire in birds (Fig. 1B). At the maximum, we identified 3,750 ORs in North Island brown kiwi (*Apteryx mantelli*), the largest number of ORs known from any animal, surpassing elephants (Fig. 1B). Conversely, we also found species with few ORs. The lowest OR counts of all birds were in crows, with only 7 in carrion crow (*Corvus corone*).

**Fig. 1.**
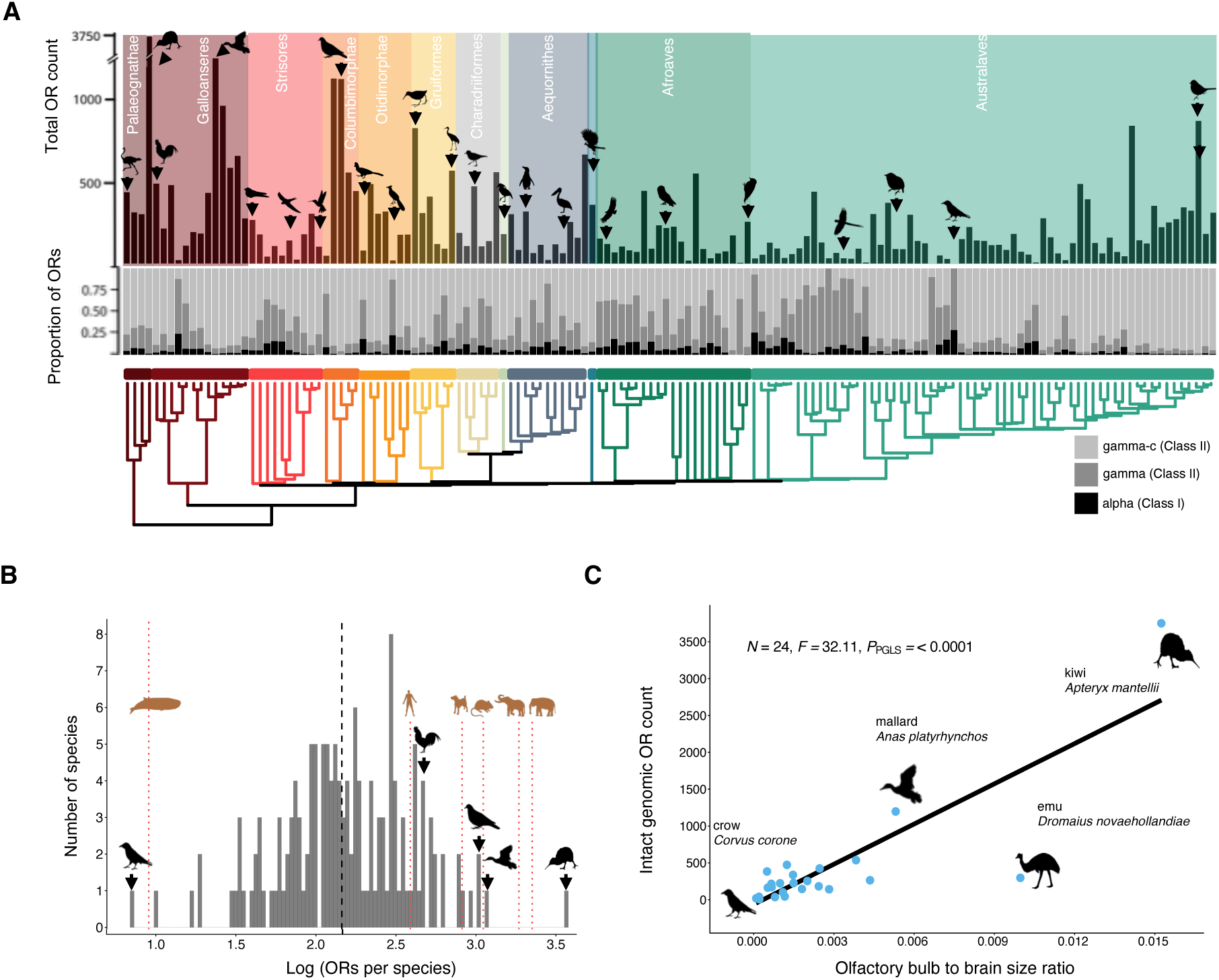
Intact genomic OR repertoires in birds. **(A)** Intact OR counts from 148 bird species assemblies. Black bars show total intact genomic OR counts. Gray bar graph shows the proportion of each species’ genomic repertoire across Class I alpha receptors (black), Class II gamma receptors (dark gray), and Class II gamma-c receptors (light gray). **(B)** Log range of total intact genomic OR counts in birds. We binned total OR counts into sets of log(0.025) ORs. Dashed black line shows the median count for birds Mammalian OR counts are shown with red lines and are from Policarpo et al.^12^ **(C)** Bird total intact genomic OR counts correlate with olfactory bulb to brain size ratio in birds. Correlation measured with phylogenetic generalized least squares test (*N* = 24, *F =* 32.11, *P*_PGLS_ *=* <0.0001).

In animals, the size of the olfactory bulb relative to the whole brain is a long-standing measurement used to assess potential reliance on the olfactory system (*17*, *18*). Following phylogenetic correction, we found a positive correlation between the revised counts of intact ORs from genomes with long-read technology (see above) with the size of the olfactory bulb to brain ratio, both overall (*N* = 24, *F =* 32.11, *P_PGLS_* < 0.0001, Fig. 3A) and individually within each subclass of OR (fig. S2). This result is consistent with bird ORs, including gamma-c, functioning in the olfactory system. Taken together, like in mammals, bird ORs are diverse and vary substantially between species likely reflecting extensive gene duplication and losses associated with the reliance on smell.

### Evolutionary Dynamics of Avian-specific Gamma-c ORs

ORs are broadly grouped into either Class I (or alpha) or Class II (or gamma) based on sequence similarity (*19*). Out of a total of 37,286 ORs across all species in birds, 3.34% included alpha ORs, while 96.61% were gamma ORs (Fig. 1A). This large proportion of gamma ORs relative to alpha ORs is consistent with OR repertoire profiles across mammalian species. Within gamma, birds have a unique expansion of ORs in a specific OR subfamily (*17*, *20*). This OR subgroup, known as gamma-c ORs, constituted 78.85% (29,413 total ORs) of the total bird ORs we identified using two of the most complete bird assemblies available, including 88.5% (419 out of 473 total) of all ORs in the chicken (Genbank accession: GCA_024206055.2, *16*, fig. S3) and 96.85% (277 out of 286 ORs total) in the telomere-to-telomere zebra finch assembly (*Taeniopygia guttata*, Genbank accession: GCF_048771995.1, fig. S3). In addition to being the most numerous bird OR group, gamma-c sequences within each species are highly similar to each other, as indicated by short branch lengths on the OR phylogeny (fig. S3) and an average of 83.83% identical nucleotide (77.78% identical amino acid) sequences across 419 chicken gamma-c ORs, compared to 44.84% in 43 gamma and 51.45% in 12 alpha ORs. Unlike other OR gene families, in which orthologs from different species generally cluster together (*21*), sequence similarity between gamma-c ORs is higher within a species than between species from different bird families (Fig. 2A and fig. S4). However, this is less clear for gamma-c OR genes from closely-related species within the same bird family, where we observe interdigitation of gamma-c genes among species (fig. S5), illustrating that fully monophyletic gamma-c OR gene clades form following millions of years of evolutionary divergence.

**Fig. 2.**
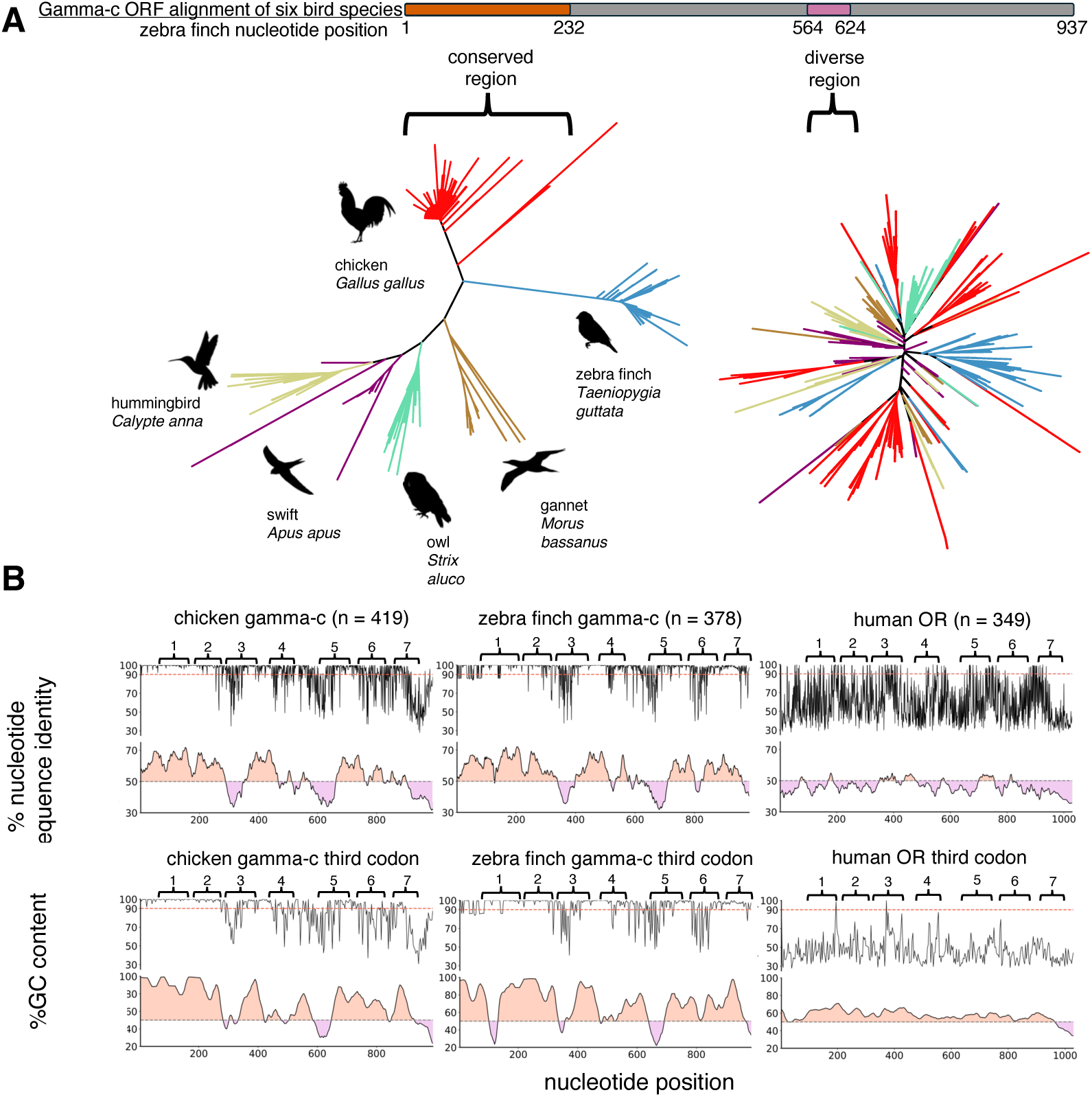
Evidence for gene conversion in bird gamma-c ORs. (**A**) Phylogenies from six bird species using subregions of gamma-c OR alignment. Gray bar shows zebra finch gamma-c ORF, with orange corresponding to a homogenized region and pink corresponding to a diverse region. Phylogeny from homogenized region (left) largely forms species-specific clades. Phylogeny from diverse region (right) forms unique phylogenetic patterns. (**B**) Regions of nucleotide sequence conservation and high GC content between ORs in bird gamma-c ORs but not in human. Aligned chicken gamma-c (left), zebra finch gamma-c (middle) and human OR repertoires (right) are shown. First row shows percent of identical nucleotides at each position in the full ORF. GC content, below sequence identity, is labelled as orange for above 50% and as lavender for below 50% GC content for 30-bp sliding window. Gamma-c OR sequence identity and GC content patterns are preserved when only observing the third codon position (second row).

The unusual species-specific phylogenetic pattern we observe between bird families (Fig. 2A) would be consistent with either parallel duplication of a single ancestral gamma-c OR in bird families or concerted evolution, including unequal crossing over and/or gene conversion (*22*). Gene conversion transfers short stretches of DNA between paralogs, typically during meiosis through DNA repair mechanisms (*23*, *24*). This process leaves several characteristic signatures (*25*, *26*): (i) incompatibility between gene trees and the true history of gene duplication, resulting in species-specific clades; (ii) the presence of subregions within genes that are nearly identical in nucleotide sequence, contrasted with other subregions that retain higher sequence diversity; (iii) a GC-bias in gene converted regions (*27*), (iv) phylogenetic relationships reconstructed from homogenized subregions versus non-converted regions are incongruent, and (v) close proximity of gene converted regions within the same chromosome.

Consistent with these predictions, we found that gamma-c ORs within species contain long stretches of highly similar nucleotides in the open reading frame (ORF). For example, in chicken (and six other bird species), an approximately 300-bp region near the 5′ end of the ORF is nearly identical within the given species across paralogs (Fig. 2B). Additional shorter stretches with high similarity occur in other parts of the ORF. Homogenized regions in chicken and zebra finch gamma-c ORs are additionally GC-rich and contrast AT-rich diverse regions (Fig. 2B). In contrast, human ORs exhibit relatively uniform diversity across the ORF and are generally AT-rich (Fig. 2B). Importantly, homogenized regions show similarity across all codon positions, including synonymous third positions, indicating that homogenization is not driven by amino acid–level selection (Fig. 2B). Statistical analyses using the program GENECONV identified gene conversion events between aligned pairwise sites in chicken gamma-c ORs (Data S1). In GENECONV, polymorphic sites between sequences pairs are assessed, with putative recombination breakpoints assigned by sequence similarity. Significant similarity across a pairwise alignment is considered recombinant by GENECONV, indicating potential gene conversion events. Our GENECONV analyses detected over thousands of recombination breakpoints in the chicken gamma-c ORs (*P* < 0.0001, Data S1) (*28*). Consistent with gene conversion events, gene trees reconstructed from homogenized versus variable regions of gamma-c ORFs were incongruent across six bird species, with significantly longer branch lengths in the latter (Fig. 2A). Moreover, gamma-c ORs are arranged in genomic clusters, often on avian dot chromosomes – for example, all chicken gamma-c ORs are found on dot chromosomes 16, 29, and 34 (*15*, *29*), and large gamma-c OR clusters are also found on specific dot chromosomes in zebra finch (*30*). Because gene conversion most frequently occurs among genes in close proximity, the persistence of chromosome-specific substitutions (fig. S6) further supports intrachromosomal gene conversion. Together, these findings indicate that gamma-c ORs are mosaics of homogenized and less-homogenized subregions, and suggest that their unusual phylogenetic pattern reflects extensive gene conversion during bird evolution.

### Expression of bird ORs in OSNs

The results above demonstrate that birds have genomic OR repertoires comparable in size to those in mammals. We next sought to determine whether and how bird ORs function in the olfactory system. To be functional and tied to olfaction, we would expect bird OR expression in the olfactory epithelium, and, specifically, the OSNs. To test this, we conducted bulk RNA-seq from olfactory mucosa and used pectoralis muscle as a negative control in the chicken, zebra finch (Genbank accession GCA_048772025.1), and brown-headed cowbird (*Molothrus ater*, Genbank accession GCA_012460135.3), and mapped reads to OR genes. We found that overall expression of ORs was over 70 times higher in the chicken olfactory mucosa compared to pectoralis muscle (Fig. 3A, Welch’s two sample t-test, *t* = 6.04, *P* < 0.001). OR expression varied among olfactory mucosa samples within a species, but was consistent with the expression levels of known OSN markers in each sample, suggesting heterogeneity in olfactory mucosa content in each sample (fig. S7). We also found that most individual OR genes were expressed in the olfactory mucosa, including the gamma-c ORs (Fig. 3B and fig. S8). Further, we found cellular-level OR expression in the chicken olfactory epithelium through *in situ* hybridization (Fig. 3C). We found that the distribution of ORs (including gamma-c ORs) in OSNs colocalized with Cnga2, an established OSN marker (Fig. 3, C and D and fig. S9). We note that, due to their high nucleotide-level similarity, our gamma-c OR probe is expected to hybridize to most, if not all, gamma-c ORs. Our observation that far more OSNs are labeled with the gamma-c OR probe than with other OR probes is consistent with this notion. This expression confirms our expectation that bird ORs contribute to the sense of smell in birds.

**Fig. 3.**
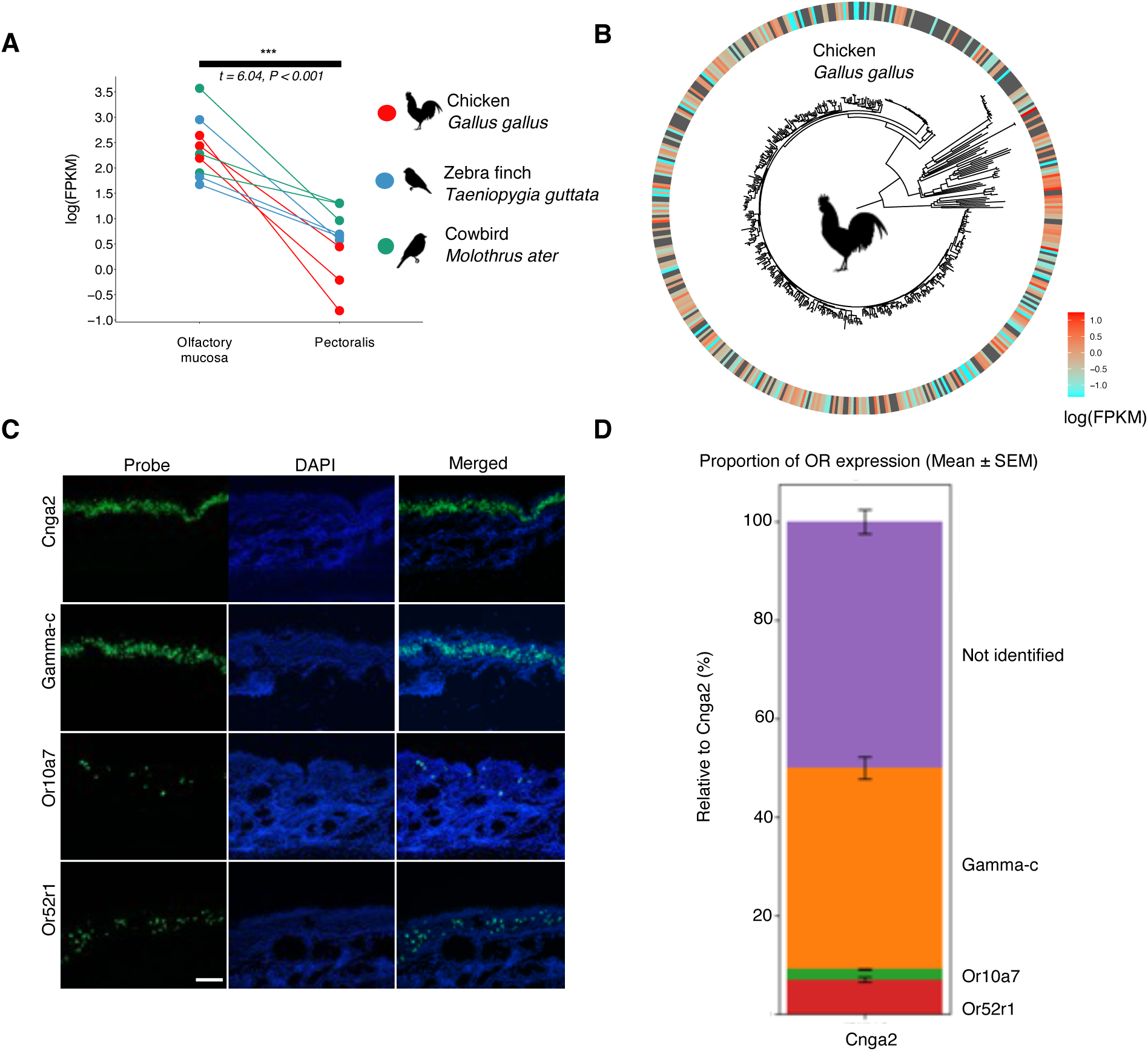
Evidence for role of OR function in bird olfactory system. **(A)** Higher levels of OR expression levels measured from olfactory mucosa and pectoralis muscle in three bird species. Significance determined with Welch’s two sample t-test (*t = 6.04, P < 0.001*). **(B)** OR expression levels of individual ORs in olfactory mucosa and pectoralis muscle tissue in chicken. Chicken is depicted with a phylogeny of its intact genomic OR repertoire. The ring shows the olfactory mucosa expression. Cyan indicates low level of expression, red high level of expression. (**C**) In situ hybridization for Cnga2, gamma-c, Or10a7, and Or52r1 (top to bottom, respectively). Scale bar shown in 100uM. (**D**) Proportions for ORs expression relative to the OSN marker Cnga2.

### Bird ORs respond to odors

Given the presence of ORs in chicken OSNs, we anticipated that bird ORs would respond to odor stimulation. First, we aimed to identify active ligands for gamma-c ORs in the chicken. Given the large number of gamma-c ORs and overall high sequence similarity, we first tested a consensus OR designed from the gamma-c OR alignment, based on our hypothesis that consensus ORs both support robust functional expression in heterologous cells and approximate the response of native ORs (*31*) (Fig. 4A, see Methods). We first tested the consensus of all chicken gamma-c ORs for a response to a panel of odors. We found that the chicken gamma-c consensus OR responded to 2-isopropyl-3-methoxypyrazine and 2-isobutyl-3-methoxypyrazine, members of the key pyrazine food odor group (Fig. 4B). Having methoxypyrazine response from a consensus OR, we next cloned native ORs directly from chicken genomic DNA (Fig. 4C). To test the function of the cloned ORs, in addition to N-terminal rhodopsin tag, we modified the C-terminal ends of each OR to enhance their expression (*32*). Several native ORs responded to 2-isobutyl-3-methoxypyrazine, 2-sec-butyl-3-methoxypyrazine, and 2-isopropyl-3-methoxypyrazine (Fig. 4D). Importantly, each OR has a unique relative response profile against a panel of pyrazines (Fig. 4D). Gene conversion mediated sequence homogenization could cause different members of gamma-c ORs within chicken to acquire similar responses to pyrazine odors, potentially limiting their ability to discriminate a broad range of environmental odors. We therefore tested additional chicken gamma-c OR responses to a variety of odors and found diverse odor responses across the tested ORs (fig. S11A). Our results show that bird ORs, particularly those within the gamma-c subfamily, have the ability to respond to odors and each OR may have unique function.

### Bird odorant receptor response to odors activating mammalian orthologs

In addition to gamma-c ORs, which are highly variable across bird species, we wanted to test odor responses from a bird OR that is stable across evolutionary time, notably, OR51E2 that belongs to the alpha (Class I) family and responds to short chain carboxylic acids. Recent studies revealed the structure and binding mechanisms of human OR51E2 (*33*, *34*). We found that chicken has an OR orthologous with human OR51E2 (Fig. 4E), and we therefore tested chicken OR51E2 response to a set of short chain carboxylic acids. We found that chicken OR51E2 responds to the same odorants, propionate and acetate, as the human ortholog, but additionally responds to butyrate, indicating moderate functional differences between the two ORs (Fig. 4F). This result demonstrates that at least one OR ortholog has conserved odor response profiles between humans and birds.

### Bird OSNs respond to odors in vivo

After demonstrating an *in vitro* response of chicken gamma-c ORs to pyrazines, we next tested for a response to pyrazines in the live chicken to determine whether OSNs expressing gamma-c ORs are activated by odor stimulation. We exposed individual chickens in a chamber to a cassette containing either 1% (v/v) or 10% 2-isobutyl-3-methoxypyrazine. We also exposed chickens to 1% acetophenone, commonly used in mammalian olfactory experiments, as well as to no-odor stimulation as controls (*35*). We then labelled gamma-c expressing neurons with *in situ* hybridization, and performed immunohistochemistry against phospho-ribosomal protein S6 (pS6) to label active neurons (*31*). In chickens receiving no odor, there was minimum pS6 labeling overlapping with gamma-c expression. Following 1% acetophenone exposure, we observed pS6-positive OSNs, but this signal did not colocalize with OSNs expressing gamma-c (Fig. 5, A and B). We found that a subset of OSNs expressing gamma-c ORs were labeled with pS6 antibodies following exposure to 1% and 10% 2-isobutyl-3-methoxypyrazine (Fig. 5A). Quantification of pS6 intensity in OSNs expressing gamma-c ORs demonstrated that pyrazine stimulation, but not acetophenone stimulation, produced a significant increase in pS6 signals in OSNs expressing gamma-c ORs relative to no odor controls (Fig. 5B and fig. S10). Together, these results support that a subset of OSNs expressing gamma-c ORs is activated *in vivo* by exposure to 2-isobutyl-3-methoxypyrazine.

**Fig. 5.**
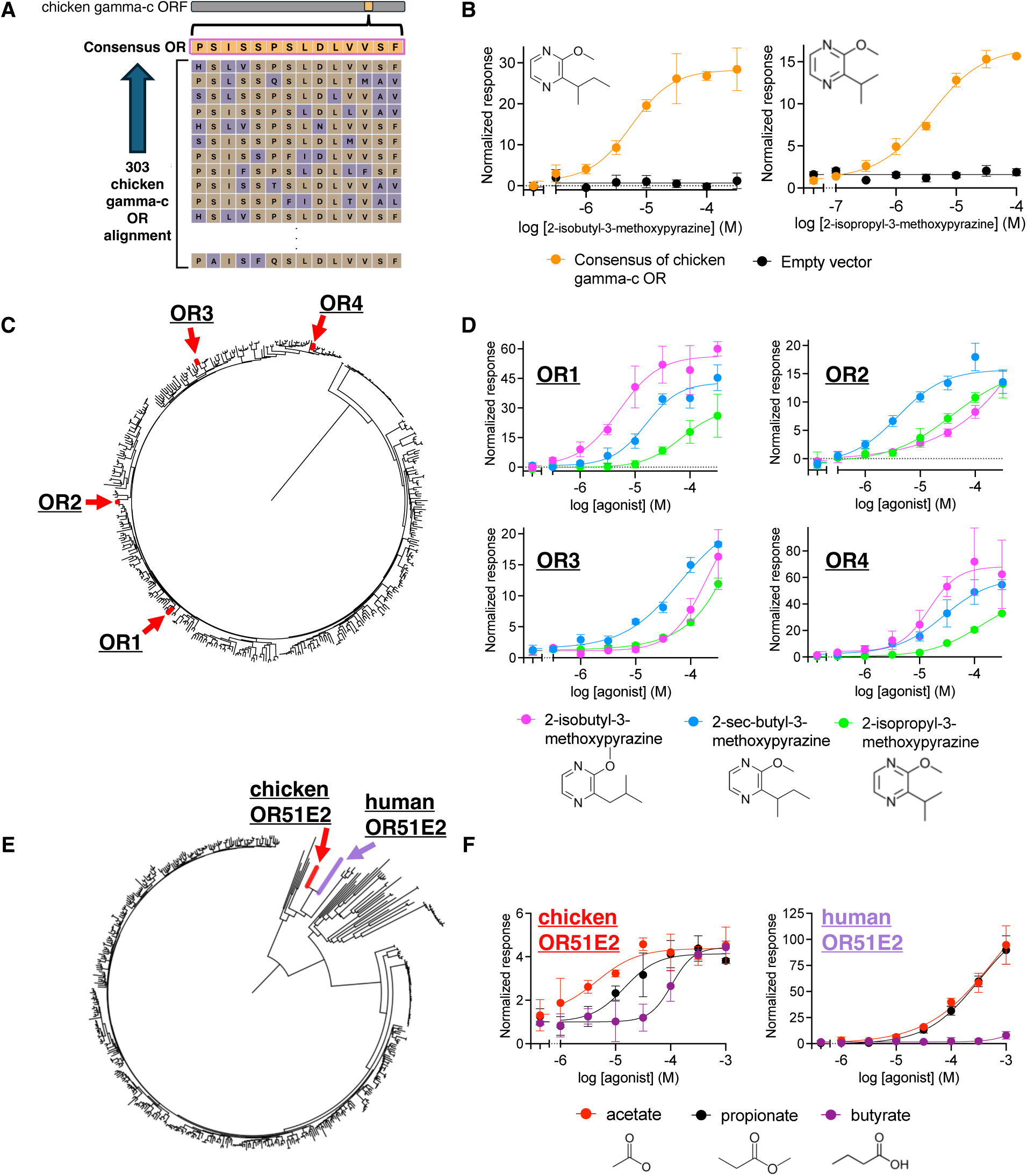
Functional testing of ORs *in vitro*. (**A**) Schematic for consensus OR sequence. Most common amino acid at each position of the gamma-c chicken OR alignment was selected to generate consensus OR sequence. (**B**) Increasing response of chicken gamma-c consensus OR (orange) with increasing concentrations of 2-isobutyl-3-methoxypyrazine (left) and 2-isopropyl-3-methoxypyrazine (right). Empty vector response is shown in black. Images are normalized to the response prior to odor addition. (**C**) Phylogeny showing the four native chicken gamma-c ORs tested against pyrazine odors *in vitro* within the chicken gamma-c OR repertoire. (**D**) Increasing response of native chicken ORs with increasing concentrations of 2-isobutyl-3-methoxypyrazine, 2-sec-butyl-3-methoxypyrazine, and 2-isopropyl-3-methoxypyrazine. (**E**) Location of human OR51E2 (red) when placed on the chicken OR phylogeny. Human OR51E2 has a 1:1 orthologous relationship with chicken OR51E2 (blue). (**F**) Response of human OR51E2 (left) to increasing concentrations of three short chain fatty acid odors, acetate (red), propionate (black), and butyrate (purple). Response of chicken OR51E2 to the same odors (right).

**Fig. 6.**
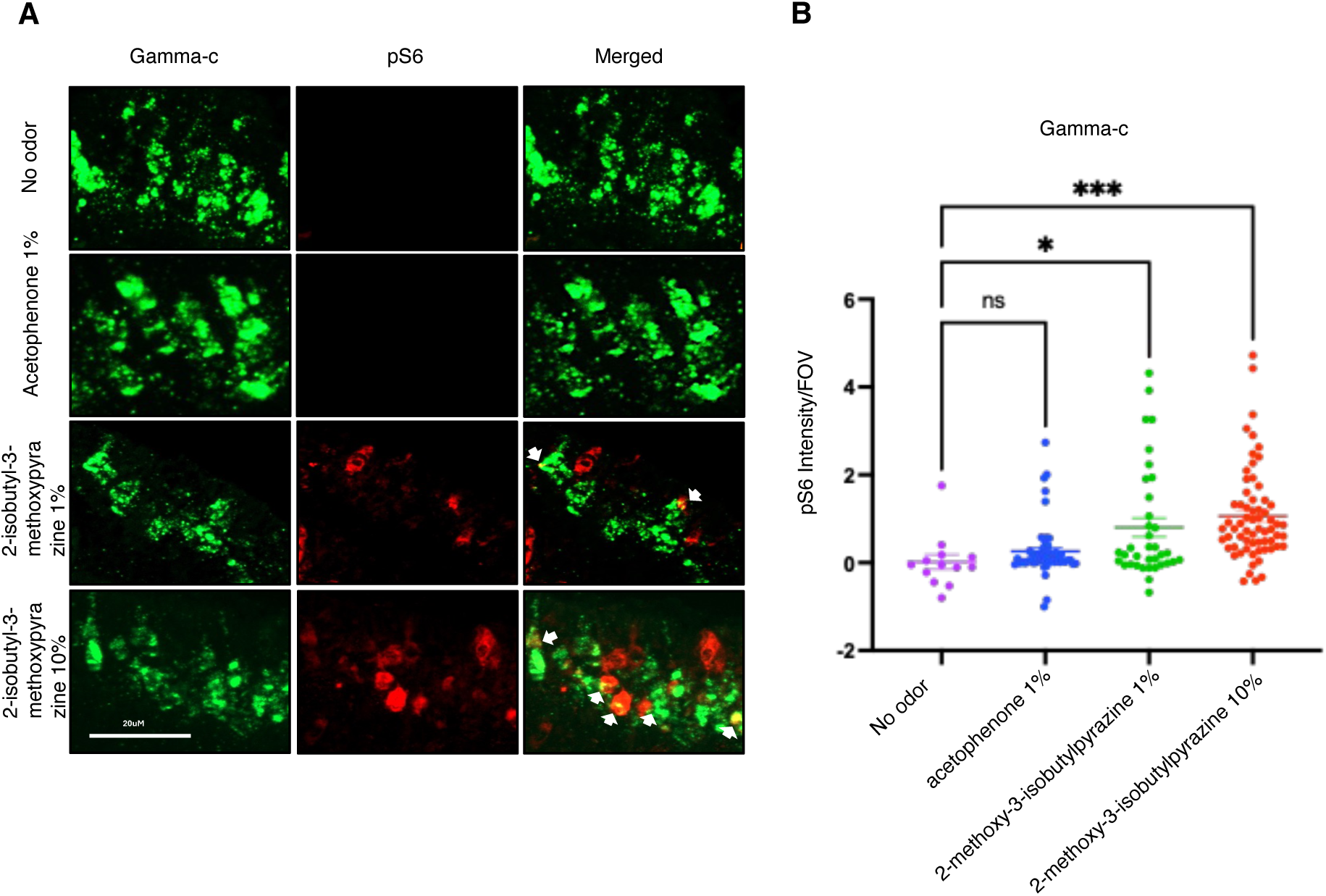
Pyrazine exposure increases pS6 in gamma-c-positive OSNs. (**A**) *In situ* hybridization combined with immunofluorescence for gamma-c and pS6, respectively. (**B**) Semi-quantitative analysis for pS6 color intensity comparing treatment groups no odor, acetophenone 1%, 2-isobutyl-3-methoxypyrazine 1% and 2-isobutyl-3-methoxypyrazine 10%. We observed significantly greater pS6 intensity in gamma-c-positive cells following pyrazine exposure compared to no odor conrol using Welch’s one-way ANOVA and Brown-Forsythe correction.

## Discussion

In this study, we show that birds often have hundreds to thousands of ORs in their genomes, and that the kiwi (*Apteryx mantelli*) has the largest OR repertoire of any vertebrate. We find that most bird OR genes belong to a bird-specific expansion of a specific Class II subgroup, known as the gamma-c ORs, and that these receptors may uniquely evolve through gene conversion. In the chicken, we demonstrate that ORs are expressed in OSNs and that some of them respond to pyrazine odors, the first odorant-receptor pairings in birds. We also show that for OR51E2, a highly conserved OR across vertebrate evolution, chicken and human ORs respond to the same short chain carboxylic acids.

### Diverse bird species have OR counts comparable to other vertebrates

We analyzed only genomes generated with long-read sequencing and a minimum contigN50 size of 7 Mb in order to avoid undercounting ORs caused by nearly identical sequences within stretches of gamma-c ORs (*15*). As a result, we find that bird OR repertoires are much larger than previously reported (*12*, *18*). For example, remarkably, the North Island brown kiwi OR repertoire increased from 63 ORs, based on the short-read genome (*12*), to 3,750 in the present study based on the long-read assembly. The kiwi forages at night and uses its bill, with nostrils located at the tip, to probe for soil invertebrates (*36*). It has the largest olfactory bulb relative to brain size of any bird, consistent with our finding that olfactory bulb size correlates with OR counts (*37*, *38*). Together, these results further support an exceptional enhancement of olfactory ability and sensitivity in North Island brown kiwi. These thousands of ORs in kiwi contrast with birds that have few genomic ORs, such as crows, which also exhibit smaller olfactory bulbs (*38*– *40*). The small OR repertoires in crows are particularly notable relative to most other terrestrial vertebrates (*12*, *13*, *41*). High variation in OR counts in birds is analogous to the pattern observed in mammals, with fewer than 20 ORs in several toothed whales but over 2,000 ORs in elephants (Fig. 1B, *12*, *13*).

Although the average OR count in birds (258 ORs) is lower than reported mean mammalian OR counts (680 ORs, (*12*)), the average is comparable to functional genomic ORs in crocodiles (276 ORs, 12), and is greater than the average OR counts reported for bony, cartilaginous, and jawless fish (*12*). As OR counts may correlate with olfactory acuity such as the ability to discriminate odor pairs (*42*), it is possible that many bird species have olfactory acuity that is similar to species with known reliance on olfaction.

### ORs are expressed in the OSNs in birds

Our study, including bulk RNA-seq in three bird species and *in situ* hybridization in chicken, suggests that the large majority of ORs, including gamma-c OR genes, are expressed in the OSNs. These results are consistent with two earlier bulk RNA-seq studies in other bird species that also showed OR expression, including gamma-c ORs, in the bird olfactory epithelium (*43*– *45*). Here we show consistent results that many genes in the alpha (Class I), gamma and gamma-c (Class II) OR repertoire also show expression in the olfactory mucosa in three species.

We further found that in the chicken, OSNs are present in the caudal turbinate of the maxilla, and that bird ORs are expressed in the OSNs. Although previous studies have shown the location of neurons in the olfactory epithelium of embryonic chicken and kiwi (*38*, *46*, *47*), this is the first time that the location of the OSNs has been shown using known OSN markers such as Cnga2 and ORs. Additionally, we saw multiple ORs that belong to alpha, gamma, and gamma-c ORs expressed in the OSNs. Within OSNs, more neurons were positive for gamma-c ORs than alpha (Or52r1) or gamma Ors (Or10a7) (Fig. 3D). Our probe used for *in situ* hybridization likely hybridizes to mRNA transcripts from most gamma-c genes due to the high overall identities and nearly identical nucleotide stretches. Therefore, we are limited in our ability to test the location of individual gamma-c ORs. This high sequence similarity of the gamma-c ORs may make it difficult to determine whether different gamma-c ORs are expressed in different OSNs, as occurs in mammals. In summary, we conclude that just as in mammals, bird ORs expressed in the OSNs of the olfactory epithelium.

### Gene conversion as an evolutionary mechanism in sensory biology

Here, we present evidence for extensive gene conversion of gamma-c ORs, resulting in highly similar paralogs within species. Such a mechanism is likely to be more common in birds, as gene conversion has also been implicated in the evolution of major histocompatibility (MHC) complex genes and toll-like receptor genes (*24*, *48*). In particular, in the chicken a large tandem cluster of the avian MHC gene family is located on dot chromosome 16, the same dot chromosome that contains some gamma-c ORs (*49*). Although the underlying reason for the apparent high prevalence of gene conversion events in birds remains an open question, this mode of evolution clearly impacts multiple gene families and shapes the evolution of genes on avian dot chromosomes.

Gene conversion could cause different members of gamma-c ORs within a species to acquire similar functional properties yet may potentially limit a species’ ability to discriminate a broad range of environmental odors. Interestingly, our data suggest that putative ligand-interacting regions of the OR, such as transmembrane domains 5 and 6, remain relatively diverse at both the nucleotide and amino acid levels (Fig. 2B). The diverse odorant responses profiles we found across chicken gamma-c ORs (fig. S11A) support the idea that gamma-c ORs within a species still retain functional diversity despite extensive homogenization via gene conversion. Additionally, prediction of chicken gamma-c OR structure using AlphaFold3 indicates that canonical binding pocket residues show higher variability (fig. S11B). Therefore, we propose an evolutionary model of partitioned gene conversion tracts that maintain structural integrity while enabling diverse ligand selectivity.

### Bird ORs respond to odors

We report an experimental response of a bird OR to an odorant by showing that chicken gamma-c ORs respond to pyrazine odors 2-isobutyl-3-methoxypyrazine, 2-sec-butyl-3-methoxypyrazine, and 2-isopropyl-3-methoxypyrazine. To humans, 2-isobutyl-3-methoxypyrazine has a potent green pepper-like aroma. In nature, pyrazines are an important class of key food odors and it is possible that chickens detect pyrazines when foraging (*50*). In one example, chickens with prolonged exposure to 2-isobutyl-3-methoxypyrazine had a larger egg size (*51*). Therefore, there could be potential implications of pyrazine odor detection and bird development. Additionally, 2-isobutyl-3-methoxypyrazine, and 2-sec-butyl-3-methoxypyrazine, 2-isopropyl-3-methoxypyrazine are characteristic odors in many insect orders, where they act as both warning, defensive, and attractant chemicals (*52–59*). Therefore, these methoxypyrazine odors may serve as important cues used by chickens during foraging and capturing insect prey.

Despite high sequence similarities, our results support the idea that different gamma-c ORs respond to different odorants based on our pS6 immunostaining showing that only a subset of gamma-c expressing OSNs responds to 2-isobutyl-3-methoxypyrazine. These results support a key role for this diverse OR subfamily in the olfactory system.

In addition to finding that odors bind to the highly evolutionary dynamic gamma-c ORs, we found that bird and human OR51E2, forming a clear orthologous relationship across many vertebrates, shows conserved response to short chain carboxylic acids. A major goal going forward will be to more completely document the functional properties of orthologs and the convergent and novel properties of paralogous ORs across taxa.

## Conclusions

Given their often bright and conspicuous coloration, as well as their heavy reliance on vocal communication, particularly birdsong, studies of birds have historically emphasized vision and audition. Our findings contribute to a growing body of work demonstrating the critical role of olfaction in avian biology. Improved genome sequencing technology, deployed with standardized protocols and annotation pipelines across the vertebrate tree of life have opened the door to a better understanding of complex gene families such as ORs. Here, they reveal that avian OR repertoires broadly overlap with, and in some cases exceed, those found in mammals. Combined with new information on the odorant-response properties of avian ORs, these results establish olfaction as an important and previously underappreciated component of avian sensory biology and provide a foundation for future studies exploring how ORs influence bird behavior, adaptation, and ecology.

## Acknowledgments

We would like to thank the Vertebrate Genomes Project (VGP) for providing the majority (76/148, 51.35%) of the high-quality long read genome assemblies used in this study. Without the VGP, we would have been unable to recover gamma-c OR repertoires from many bird species, and we greatly thank this tremendous resource. We would like to thank Dr. Kenneth Anderson, Becca Wysocky, and Christina Sigmon at North Carolina State University Department of Poultry Science for their dedication and assistance with obtaining chicken tissue and their help with chicken odor exposure experiments. We would like to thank Dr. Richard Mooney, Dr. Audrey Mercer, and Michael Booze at Duke University School of Medicine Department of Neurobiology for assistance with zebra finch tissues. We thank Dr. Marc Schmidt at University of Pennsylvania for graciously collecting brown-headed cowbird tissue. We thank Emily Xu for chicken tissue sectioning and manuscript review.

## Funding

National Science Foundation award IOS 1655730 (CNB),

National Science Foundation award DEB 1457541 (CNB)

National Science Foundation award DBI 2208965 (RJD)

National Institutes of Health grant R01DC022770 (HM)

National Institutes of Health grant R01DC020353 (HM)

National Institues of Health grant R01DC021585 (HM)

Society for Integrative & Comparative Biology (RJD)

American Museum of Natural History (RJD)

Society for the Study of Evolution (RJD)

American Genetics Association (RJD)

American Ornithological Society (RJD)

Wilson Ornithological Society (RJD)

## Author contributions

Conceptualization: RJD, CNB, HM

Methodology: RJD, MAM, VJK, PM, WS, RJL, MSB, HYL, KFPB, MS, GF, IO, NEM, HM, CNB

Investigation: RJD, MAM, VJK, PM, WS, RJL, MSB, HYL, KFPB, MS, GF, IO, NEM, HM, CNB

Visualization: RJD, MAM, VJK, WS, RJL, HM, CNB

Funding acquisition: RJD, HM, CNB

Project administration: RJD, HM, CNB

Supervision: RJD, MAM, HM, CNB

Writing – original draft: RJD, HM, CNB

Writing – review & editing: RJD, MAM, VJK, PM, WS, RJL, MSB, HYL, KFPB, MS, GF, IO, NEM, HM, CNB

## Data, code, and materials availability

Data that support the findings of this study are available in Dryad with the identifier DOI: 10.5061/dryad.0zpc867b4. All RNA-seq files are in the process of uploading to NCBI and are available upon author request. The authors declare that all data supporting the findings of this study are <u>available</u> within the paper and its supplementary files.

## Supplementary Materials

Materials and Methods

Figs. S1 to S11

References (*60*–*87*)

Data S1 to S#

