## Supplemental materials for "Unique evolutionary radiation of odorant receptors in birds"

Science
AAAS

### Supplementary Materials for

#### Unique evolutionary radiation of odorant receptors in birds

Robert J. Driver[1,2*], Mona A. Marie[2], Victoria J. Ko[2,†], Priyanka Meesa[2,‡], Wanting Sun[2], Renee J. Li[2], Michael S. Brewer[1], Hsiu-Yi Lu[2], Kevin F. P. Bennett[3], Marco Sollitto[4,5], Giulio Formenti[5], Ichie Ojiro[2,6,§], Nivritti E. Mantha[2,7], Hiroaki Matsunami[2,8*], and Christopher N. Balakrishnan[1,9,10]

**The PDF file includes:**

Materials and Methods
Figs. S1 to S11
Tables S1 to S9

**Other Supplementary Materials for this manuscript include the following:**

Data S1 to S25

#### Materials and Methods

##### In vivo *animal studies*

The RNA sequencing study design looked for the presence and expression levels of ORs in bird olfactory mucosa and pectoralis muscle. Five individuals of four species were obtained in this study for gene expression analyses: *Gallus gallus*, *Taeniopygia guttata*, and *Molothrus ater*. In all three species, individuals were obtained from captive group-housed populations and all individuals were in good health. Individuals were sacrificed as part of ongoing experiments in their laboratory. Obtaining olfactory mucosa and pectoralis tissue was a byproduct of previously scheduled sacrifices for other purposes. IACUC protocols for proper care and sacrifice were followed for the associated ongoing experiment. For *Taeniopygia guttata*, and *Molothrus ater*, all individuals were wild-type adults. For *Gallus gallus*, individuals were 21-week-old Hyline W-36 white leghorns. For *Molothrus ater*, all indiviuals were male. For *Gallus gallus* and *Taeniopygia guttata*, all individuals were female.

For *in situ* hybridization, we used three female *Gallus gallus* Hyline W-36 white leghorns at 60 weeks of age.

##### In vitro *cell lines*

All cell-based assays used Hana3A cells, a cell line derived from HEK293T cells, originally from an immortalized human embryonic kidney cell lines. Hana3A cells express RTP1L, RTP2, REEP1, and $G_{aolf}$, proteins that aid in the expression of ORs and the transport of ORs to the cell surface (*60*). We grew Hana3A cells in Gibco Minimal Essential Media ThermoFisher FBS, 5mL Gibco Glutamax, Gibco streptomycin, and penicillin.

Assembly selection

We investigated OR diversity in birds by selecting publicly available genome assemblies on GenBank (Data S2, https://www.ncbi.nlm.nih.gov/genbank/). Assemblies for each species implemented some form of long-read sequencing technology, including Pacific Biosciences or Oxford Nanopore methods. Genomes varied in the assembly methods used and in the size and total number of contigs and scaffolds. We selected only assemblies using long read sequencing, with a minimum contigN50 size of 7 Mb, due to the difficulty in recovering total OR counts in assemblies with shorter contigs (*15*). In total, we analyzed 148 different bird assemblies, including species from the three main lineages of birds, the Palaeognathae, Galloanserae, and Neoaves. The species set represents diverse ecology, diets, and trophic levels.

OR identification and classification

To detect putatively functional ORs in the selected genomes, we created a BLAST query with a set of 2,110 OR protein sequences from 6 mammals (*Ornithorhynchus anatinus, Didelphis virginiana, Bos taurus, Canis lupus, Rattus norvegicus, Macaca mulatta*), 2 birds (*Gallus gallus, Taeniopygia guttata*), and 1 crocodilian (*Gavialis gangeticus*). We obtained this query OR set by combining previously published OR subgenomes (*17*, *20*, *41*). Using this query file, we performed TBLASTN searches against all 148 bird genomes with a threshold of $E < 1\text{e}{-20}$. The TBLASTN –num_alignments option was set to 200,000 to capture all genomic ORs similar to a single query sequence. To remove pseudogenized and truncated ORs, we filtered for hits > 250 amino acids long. For any single location on the genome, we filtered out hits within 100 bp of each other, and selected the lowest *E*-value associated with that location.

After obtaining unique BLAST hits, we extracted the associated nucleotide sequence from the genome as well as 300-bp regions flanking the hit both upstream and downstream. We used a modified Perl script to detect open reading frames (ORFs) within each extracted region (*53*, *54*). We then aligned these ORFs to each other as well as to the human Olfactory Receptor Family 2 Subfamily J Member 3 (OR2J3) sequence using the E-INS-I default parameters in MAFFT (*63*). Using the previously characterized transmembrane domains of OR2J3 as a guide, we removed any sequences that had five or more amino acid insertions or deletions within a transmembrane domain in the alignment (*62*, *63*). This included ORFs with stop codons appearing prior to the end of the seventh transmembrane domain. We performed this filtration with custom python scripts (Data S3).

Using this alignment, we recorded the position of the first amino acid in the first transmembrane domain. To estimate the location of the ORF start codon, we used modified Perl scripts to find the most appropriate methionine upstream of this recorded transmembrane start position (*53*, *54*). ORF sequences were then truncated at the 5' ends to begin with this methionine. This set of ORFs was then aligned using the E-INSI-I parameters in MAFFT (*55*) to a set of *T. guttata* reference ORs as well as 11 non-OR rhodopsin-like G-protein coupled receptors (non-OR GPCRs) that functioned as an outgroup (*62*). We then used clustalW to generate a neighbor-joining tree from this alignment with 1000 bootstraps, gaps removed, and Kimura's distance correction (*64*, *65*). We then removed any ORFs that were phylogenetically more closely related to the non- OR GPCRs.

We classified all remaining ORFs as functional ORs. Using this final set, we ran a maximum likelihood tree using IQ-TREE with automatic model selection and 1000 SH-like approximate likelihood ratio test replicates (*58*). Using ML support values, we collapsed all nodes <50%

support into a polytomy using iTOL software, and rooted the tree using the ancestral branch leading to the 11 non-OR GPCRs (*59*). We classified bird ORs into subfamilies alpha, gamma, and gamma-c based on the subfamily of the query sequence used to identify the OR and the location of the OR in one of the three distinct avian OR clades (*14*, *41*). We then counted the final number of OR sequences as well as the number of ORs from each subfamily.

<u>*Estimation of tree topology*</u>

To analyze OR counts in a phylogenetic context, we sought to create a phylogeny of the surveyed bird species. The bird species used in this study are a unique set, with no preexisting published phylogenies containing the species in a single tree. Therefore, we used topologies from existing phylogenies in the literature. We used an established bird phylogenetic tree as the topology for our tree structure (*60*). Three species in our analysis were not present in the original phylogeny- *Manacus candei, Pyrocephalus nanus,* and *Oenanthe melanoleuca*. These species are a result of phylogenetic splits, but the sister species were present in the original phylogeny (*60*). Therefore, the species in the dataset were placed at the position of their respective sister species in tree (*60*). We then used the drop.tip () function with the *ape* package (*61*) in R to remove all species not present in our analysis, leaving us with a tree of species with OR counts. Our resulting tree had polytomies in the tanager group, which we then resolved manually using reference topologies.

<u>*Estimation of branch lengths*</u>

To determine the branch lengths for our literature-based topologies, we mined the genomes of our 148 species subset for ultraconserved elements (UCEs). We downloaded fasta files from GenBank (accession numbers of assemblies in Data S2). We next followed the UCE discovery procedure recommended in PHYLUCE (*70*) to extract UCE loci from reference genomes. We next aligned UCE loci from all genomes using MAFFT (*55*) and trimmed using GBLOCKS (*71*), both implemented in PHYLUCE. We subset the alignments to retain only those with no missing data and concatenated them into a single alignment. We ran IQ-TREE (*58*) using a subset of 15 taxa and the model finder option to select a sequence evolution model. We ran IQ-TREE on the full dataset constrained to the topology of Stiller et al. (*64*) and using the best model from the subset run (TVM+F+R6). Finally, with R package ape (R Core Team 2024, *61*) we converted the resulting tree to an ultrametric tree using the relevant calibration dates used by Stiller et al. (*64*).

*Percent identity and percent GC content plot generation*

Alignment of all chicken gamma-c OR, zebra finch OR, and human OR was generated and used to create a third codon aware alignment. We first aligned protein sequences with MAFFT, then back-translated the protein alignment into a codon alignment using PAL2NAL (*65*). We obtained the human OR sequences through the HORDE database (*74*). Percent identity per nucleotide position was calculated by first filtering each individual alignment, removing positions >95% gaps, generating consensus sequence, and using the following equation:

$$\%\ \text{identity per nucl position} = \frac{\#\ \text{nucl the same as consensus at given position}}{\text{Total}\ \#\ \text{nucl (excluding gaps)}}\ \text{x}\ 100\%$$

Percent GC content plot was calculated using a sliding window approach. For each 30-position sliding window bin, we calculated GC content by:

$$\%\ \text{GC content} = \frac{\#\ \text{G} + \text{C nucl (excluding gaps)}}{\text{Total nucl in window (excluding gaps)}}\ \text{x}\ 100\%$$

*Olfactory bulb size: phylogenetic generalized least squares*

The olfactory bulb size relative to the brain size was available for 24 species in our dataset, published in Corfield et al. (*37*). We omitted species in this analysis that were not represented in the published dataset. To control for the phylogenetic non-independence of our trait comparisons across bird species, we ran phylogenetic generalized least squares (PGLS) models. The phylogenetic trees with branch lengths generated from the UCE dataset were converted to a correlation structure in R using the *ape* package function corBrownian to estimate a Brownian motion (BM) model of trait evolution and corMartens to estimate an Ornstein-Uhlenbeck (OU) model (*67*). The OU model may better replicate actual biological processes due to an additional parameter to the "random walk" of BM in that there is a greater attraction to an initial central value the further the trait is from this value. We then used the function gls in the R *nlme* package. For each trait comparison, we compared the AIC values of each model to determine whether to select BM or the additional parameter in OU. We then ran ANOVA tests on our models followed by general linear hypotheses tests to determine significance, using the *multcomp* package in R.

*Olfactory mucosa sample collection (for downstream RNA-seq)*

To determine the location of the bird mucosa and specific mucosa regions (the anterior, middle, and posterior conchae), we referenced morphological descriptions and images of the maxilla. We originally practiced dissections on bird carcasses donated by the North Carolina Museum of Natural Sciences. In this unique dissection, the maxilla was cut transversely through the nares and then from this incision the sides of the maxilla were cut proximally towards the lores. There were three cuts in the maxilla, one transverse and distal, the other two sagittal from the nares to the lores. From this, the proximal half of the maxilla can be lifted up from the nares, exposing the tissue in the maxilla. We sampled as much tissue as possible in this part of the maxilla, and tried to sample from all three regions (anterior, middle, posterior) of the conchae, and placed immediately in microcentrifuge tubes on dry ice. Following sample collection, samples were stored in -80 C freezers. We obtained pectoralis muscle tissue at the same time, following olfactory mucosa sampling.

We obtained olfactory mucosa from three bird species: *Gallus gallus (*chicken), *Taeniopygia guttata (*zebra finch), and *Molothrus ater* (cowbird). In total, we obtained three olfactory mucosa and pectoralis samples from each species. R.J.D. sampled the chickens immediately following a routine dispatch in the laboratory of Dr. Ken Anderson at the Prestage Department of Poultry Science at North Carolina State University. The chickens were 21-week old hyline W-36 white leghorn hens (female). R.J.D. collected the zebra finch samples from the laboratory of Dr. Richard Mooney in the Department of Neurobiology at the Duke University School of Medicine. All zebra finches were adult females from separate parents. Dr. Marc Schmidt at the Department of Biology at the University of Pennsylvania collected and dissected the cowbirds. All brown-headed cowbirds were adult males. All four species were sampled from captive populations, including the domesticated chicken and zebra finch.

*RNA extractions and sequencing*

To extract RNA from the olfactory mucosa and pectoralis tissue, we cut a small amount of tissue (roughly 2x2 cm) from each sample, and cut samples on dry ice. We immediately transferred tissue to 1mL RNAzol RT (Molecular Research Center, Inc., Cincinnati, OH) according to the manufacturer's brochure (March 2017), and dissolved the sample with a homogenizer. We then added 400uL water to DNA, protein, and polysaccharides, and then waited 15 minutes to precipitate. We centrifuged to remove these at 12,000 g for 15 minutes. We next added 5uL 4-bromoanisole to 1mL of supernatant for phase separation, waited 3-4 minutes, and then centrifuged at 12,000 g for 10 minutes. We performed this optional step of the protocol twice. We then precipitated the isolated RNA by adding equal volume isopropanol to the supernatant, waited 15 minutes, and then centrifuged 12,000 g for 10 minutes. We then washed with 400uL 75% ethanol and spun at 4,000 g for 3 minutes, and repeated this step twice. We then solubilized in water. We tested RNA concentration and purity using a NanoDrop spectrophotometer (Thermo Fisher Scientific, Waltham, MA, USA), and RNA quality and integrity were assessed with an Agilent 2100 Bioanalyzer (Agilent Technologies, Santa Clara, CA, USA) at the Brody Integrative Genomics Core in the Department of Pathology & Laboratory medicine at East Carolina University.

We examined RNA quality with the 4200 TapeStation (Agilent Technologies, Santa Clara, CA), with RNA integrity number (RIN) of samples ranged from 6 to 10. We determined RNA concentration with the Qubit Fluorometric Quantitation (Thermo Fisher, Waltham, MA), with 150 ng of RNA samples used for each NGS library preparation. We prepared stranded cDNA libraries using the TruSeq Stranded LT mRNA kit (Illumina, San Diego, CA) in accordance with the manufacturer's protocol using the poly-adenylated RNA isolation. We performed sequencing of paired-end reads (100 bp × 2) by pooling all the samples together on the NextSeq 2000 system with a P3 200 cycles reagent. We de-multiplexed and trimmed raw sequence reads for adapters the on-instrument DRAGEN GenerateFastQ pipeline (v3.7.4).

*Read mapping*

We mapped reads using the Spliced Transcripts Alignment to a Reference (STAR) aligner (*76*). We were interested in OR expression specifically, so we generated the STAR reference genome not from the available species genome assemblies, but from our previously established genomic

OR repertoires of each species. Additionally, not all ORs are annotated in previously published assemblies. We found the genomic OR repertoires for chicken, zebra finch, and cowbird from our previously described genomic scans. From our final curated OR alignments, we used custom R scripts and bedtools to extract nucleotides from the associated genome. We generated the reference genome of OR sequences without using a GTF reference annotation. We then mapped reads to the genomic OR repertoires using STAR default parameters. We also mapped all other genes back to each species reference genome to find the expression of all genes in the olfactory mucosa and pectoralis samples.

*Counting and differential expression*

We counted the number of reads in output SAM files using the *dplyr* package in R (*77*). To measure gene expression, we converted raw counts to fragments per kilobase of transcript per million (FPKM). We did not filter genes with low expression due to previous reports of many bird ORs showing low expression levels. We used a standard linear model with "tissue" (either pectoralis or olfactory mucosa) as the independent variable testing within *Gallus gallus*, *Taeniopygia guttata*, and *Molothrus ater*. We ran a Welch's two sample t-test comparing log-transformed FPKM values between olfactory mucosa and pectoralis samples for chicken, zebra finch, and cowbird, as an alternative way to measure differential expression from a relatively small number of overall genes. For mapping to phylogenetic trees, we used trees created as described previously, using maximum likelihood methods in IQ-TREE (*58*). We overlayed expression heatmap plots to the phylogeny using the gheatmap function in *ggtree* in R (*70*).

*Olfactory epithelium sectioning*

We used three female *Gallus gallus* Hyline W-36 white leghorns at 60 weeks of age. Following sacrifice and decapitation, we transported chicken heads on ice and dissected the chicken by cutting the maxilla transversely and collecting tissue within. We obtained chicken turbinate tissue at the posterior or proximal end of the maxilla. We then embedded the dissected turbinate in optimal cutting temperature (OCT) compound histology mold. We stored embedded tissue in liquid nitrogen before transferring to -80 C. We then used a Leica CM 1850 Cryostat to section 18um tissue sections onto VWR superfrost frosted adhesion slides. We stored slides at -80 C.

*OR probe preparation: PCR amplification of DNA template*

Using chicken genomic DNA as template, we followed steps outlined previously (*79*). We set up a PCR reaction (Table S1) using primers designed to amplify chicken ORs from the alpha, gamma, and gamma-c subfamilies (Data S4) and ran for 25 cycles (Table S2). We then cleaned up amplified DNA template using the MinElute columns in the Qiagen MinElute PCR purification kit [Qiagen #28004], which yields a final elution in 10uL of EB buffer (10 mM Tris-HCl, pH 8.5). For purification, we first added 200uL PB buffer to the PCR reaction and mix thoroughly. Then, we transferred the mixture to MinElute columns and centrifuged at full speed for 30 seconds. We then washed columns with 750uL PE buffer, spun for 30 seconds at maximum speed in a microcentrifuge, and discarded flow-through. Finally, we centrifuged for two minutes at maximum speed to remove any residual PE buffer. We then transferred the MinElute column to a 1.5mL microcentrifuge tube. We then added 10uL of elution buffer directly to the center of the MinElute column and centrifuged for 2 minutes to elute the PCR product. We then loaded 1uL of the purified DNA template on a 1% agarose gel and ran gel electrophoresis to check for the recovery of appropriately sized PCR product. To be permissible, a strong and distinct band is necessary.

*Probe synthesis, hydrolysis, and cleanup*

We then set up the transcription reaction (Table S3), and incubated for 120 minutes at 37C. We then prepared alkaline buffer (80mM $NaHCO_3$, 120mM $Na_2CO_3$), making 150uL total, or 12uL of 1M $NaHCO_3$, 18uL of 1 M $Na_2CO_3$, in 120uL of nuclease-free water. We then added 25uL alkaline buffer to the transcription reaction and incubated at 60C for 15 minutes. We then purified the reaction using RNase-free riboprobe purification columns in the Micro Bio-spin 30 chromatography column (Cat #732-6223, Bio-Rad Laboratories, Inc., USA). Prior to using the columns, we inverted columns multiple times to mix the Bio-gel resin and remove bubbles. We then removed the column's bottom tip and place in a collection tube. We next spun down the resin in the column at 3400rpm for 2 minutes and then place the column into a 1.5mL microcentrifuge tube. We pipetted the RNA probe solution directly onto the resin and then centrifuged at 3400rpm for 4 minutes. Following elution, we added 35uL UltraPure Distilled Formamide (Cat #15515-026, Life Technologies Corp., CA, USA) to the probe. We then ran 5uL of the eluted probe in a 1% agarose gel electrophoresis. The correct sized products appeared as a fuzzy band between 100 and 200bp. We then stored probes at -80C.

*RNA fluorescent in-situ hybridization and immunofluorescent Staining*

We first removed the slides with sections from the -80 C and placed on a clean flat plastic tray cleaned with 70% ethanol. We then took a hairdryer and blow-dried the slides until dry. We then loaded on a slide rack and dipped the slides in a staining bucket of 4% Paraformaldehyde (Cat#S898-07, Avantor Performance Materials LLC., PA, USA) for 15 minutes. Next we washed slides in a bucket of 1X PBS for 5 minutes, and repeated this step for a total of two washes. Meanwhile, we prepared a triethanolamine solution in a 1L glass beaker, gently mixing 8.2mL

Triethanolamine (Cat # 9468-01, Baker Analyzed, Avantor Performance Materials, Inc., PA, USA) with 800mL dH2O using a magnetic stir bar. We dipped the slide rack into the solution while the stir bar was spinning. We submerged the slides for 10 minutes, and during the first minute, we added 1.75mL Acetic Anhydride (Cat# A-6404, Sigma Chemical Co., MO, USA) dropwise while stirring. Following this submersion, we then wash the slide rack in a new bucket of 1X PBS for 5 minutes.

We then set up slide holders using a 150mm diameter Petri dish with blotting paper on the bottom, and two 1mL serological pipettes placed on top for holding slides. We soak the blotting paper in 5x saline sodium citrate buffer (SSC) (Cat # AB131156, American Bioanalytical, MS, USA) with 50% UltraPure Distilled Formamide (Cat #15515-026, Life Technologies Corp., CA, USA). We then dried the slides using blotting paper and placed the slides on the slide holders. We then added 500uL of prehybridization buffer and incubated for at least one hour at 58C. We then added the DIG probe to new prehybridization buffer at a concentration of 1uL per 200uL of buffer and heat at 85C for 5 minutes. We then added 200uL of the probe and prehybridization buffer mix (50% formamide (Invitrogen 15515-026), 5xSSC (American Bioanalytical AB13156), Baker's yeast RNA (Sigma R-6750, 250ug/ml), Herring or Salmon sperm DNA (Sigma D7290 or D1626, respectively, P/C 100ug/ml), 1mM DTT, heparin (Sigma H3393, 300U/ml)) to each slide in the slide holder. We covered the slides with strips of parafilm to form a coverslip and incubated overnight at 58C.

The next day, we warmed 5X Saline Sodium Citrate buffer (SSC) (Cat # AB13156, American Bioanalytical, MA, USA) and 0.2X SSC to 72$^{o}$C in an incubator. We then made a 5% blocking solution stock, of 10g blocking reagent (Cat #11096176001, ROCHE Diagnostics GmbH, Mannheim, Germany) in 200mL maleic acid buffer and stored at 4C. We then diluted this solution to 0.5% blocking solution in 1X PBS, and dilute 7.5% Bovine Serum Albumin (BSA) stock (Cat # 15260-037, Gibco, Thermo Fischer Scientific, MA, USA) to a 0.1% BSA solution in 1X PBS. We prepared a working solution of DIG-POD antibody by making a 1:1000 dilution of Anti-Digoxigenin-POD, Fab fragments (Cat# 11207733910, ROCHE Giagnostics GmbH, Mannheim, Germany), in the 0.5% blocking solution.

We filled two buckets with the 5X SSC and two with 0.2X SSC. We then took the slides and dry with blotting paper, and dip slides into the warm 5X SSC. The parafilm coverslip should fall off in the buffer, but if it did not, we removed with forceps. We then transferred the slides to a second bucket of 5X SSC and then moved to a bucket with the warm 0.2X SSC at 72C for 30 minutes. We repeated this step in a second bucket of 0.2X SSC at 72C for 30 minutes. We then moved slides to a wash step with a bucket of 1X PBS for 5 minutes at room temperature. We then moved slides from the slide rack to a slide mailer containing 0.5% blocking solution and incubate for 30 minutes. We then removed slides from the mailer and use blotting paper to removed excess solution. We then placed each slide back in the slide holder, with 300uL of antibody placed on top of each slide. We incubated the slide holders 45 minutes at room temperature. We then removed the antibody solution with blotting paper, placed slides back in mailers, and rinse twice with 1X PBS. Following rinses, we then washed with 1X PBS in the slide mailer three times, each for 10 minutes. We then placed the slides in 0.1% BSA solution and prepared TSA working solution. The recipe for TSA working solution is 0.003% $H_2O_2$ and a 1:400 dilution of FITC-tyramide stock in in 1X PBS. The FITC-tyramide is prepared following previous protocols (*80*, *81*). Tyramide-FITC was generated using fluorescein-NHS ester (Cat # 46100, Pierce), tyramide (Cat # T-2879, Sigma), Dimethyl formamide (DMF) (Cat # T-8654), and triethyl amine (TEA) ( Cat # T-0886, Sigma), prepared by by mixing 4 ml FITC NHS in

DMF and 1.37 ml tyramide solution and incubated in the dark in RT for 2 hrs, then added 4.6 ml ethanol. We diluted the H2O2 from an original 30% H2O2 stock in 1X PBS. After this, we removed slides from the slide mailer and removed excess fluid using blotting paper. We then add 300uL of the TSA working solution per slide and incubate for 10 minutes at room temperature in the dark. We then carefully TSA working solution using blotting paper and placed slides back to in the slide mailer. We then rinsed slides twice with 1X PBS and then wash with 1X PBS twice for 5 minutes each. If only FISH is required, we moved to the counterstaining step. If double labeling with phosphorylated S6 (pS6) antibody is required with continue with Immunofluorescent (IF) staining.

For double-labeling with the pS6 antibody, we blocked FISH labeled sections (0.1% Triton-X, 5% skim milk, in PBS) for 30 minutes, then incubated with anti-phospho-S6 (244/247) 1:300 (Cat # 44-923G, Thermo Fisher Scientific, MA, USA) diluted in blocking solution overnight at 4°C. After washing with PBS, we incubated sections with the donkey Cy3-conjugated anti-rabbit IgG 1:200 (Jackson ImmunoResearch Laboratories, PA, USA) for 45 min, washed again, and counterstained the nuclei.

For counterstaining, we replaced the 1X PBS with nuclei staining solution. To make the staining solution, we made a solution of 1% or 25uL Hoechst nuclear stain (bisbenzimide H33258, Cat # B2883, SIGMA-ALDRICH Co. St. Louis, USA), in 250mL 1X PBS, and incubated in the dark for 5 minutes. We then discard the nuclei staining solution and washed slides twice with 1X PBS for 5 minutes. We then have a final rinse with dH2O. We finally dried the excess liquid using blotting paper, and then we sealed the slides with Mowiol mounting media and a coverslip, and stored them in a slide box at 4°C.

*Imaging for in situ hybridization and pS6*

Images were acquired as z-stacks of optical sections at 1 μm intervals using a Zeiss Axio Observer Z1 inverted microscope equipped with a Zeiss Axiocam MRm camera at 200× or 400× magnification. The orthogonal projection function was used to generate composite images for visualization and quantification purposes.

*Quantification of pS6 images*

For quantification of percent of probe-positive cells, ImageJ2, version 2.9.0/1.53t was used. We quantified the total cell number per field of view (FOV) in the blue channel, and the probe-positive cells in the green channel, to calculate the percent of expression of probe positive divided by total cells counted in the FOV. Data processing and statistical analysis were performed in Python (version 3.11) using standard scientific libraries including NumPy, Pandas, and Matplotlib. The mean and standard error of the mean (SEM) were calculated for each experimental group using SEM=SD/√n, where SD is the standard deviation and n is the number of biological replicates. Graphical visualization of the quantified ISH data, including bar plots with mean ± SEM, was generated using Matplotlib. The plotting code was custom written in Python and applied uniformly to all datasets to ensure consistent normalization and scaling.
For pS6 signal quantification, the pS6 pixel intensity was quantified in individual cells labeled with the mRNA probe (OR-expressing) using ImageJ2, version 2.9.0/1.53t, and normalized values were analyzed. Statistical comparisons were performed using Welch's one-way ANOVA (Brown–Forsythe correction) to account for unequal variances and sample sizes, followed by Dunnett's multiple comparisons test to compare each condition to the control group (532).

*Design of consensus OR sequence*

To screen for many ORs at once initially, we designed consensus OR sequences representing the sequences of multiple ORs. For example, in the chicken, we took the amino acid sequences of all 303 chicken gamma-c ORs (*15*) recovered from the genomic repertoire of the genomic repertoire of chicken (NCBI accession GCF_000002315.6) and aligned these ORs using the E-INS-I default parameters in MAFFT (*55*). We then selected the most common amino acid residue at each position in the alignment, and created a "consensus OR" of all of the most common residues from the 5' to the 3' end of the OR ORF. The stop codon was assigned at the first consensus stop codon in the alignment. In the infrequent case that multiple residues were equally common at a single position in the alignment, we picked the amino acid at random. We trimmed all alignments at the 5' end prior to the first consensus methionine, and trimmed all alignments at the 3' end following the first consensus stop codon. We removed all residues corresponding to an insertion that was not present in the majority of ORs, or a deletion that was present in the majority of ORs.

***OR cloning procedure***

We designed primers for OR amplification (Data S5) based on the OR 5' and 3' sequence. We designed primers to have an estimated denaturation temperature of 56 C to 58 C, or roughly 18-22 nucleotides from the 5' or 3' end of the OR. We then added a 5' forward MluI restriction enzyme linker *AAACGCGT) and a 3' reverse NotI linker (TTGCGGCCGC).  We dissolved primers in Gibco water to 100uM and then made a 5uM working stock. For OR consensus sequences, we first centrifuged the tube, then added 100uL of TE buffer to reach a final concentration of 10 ng/uL. We then vortexed and incubated at 50 C for 20 minutes. We then created a PCR mix (Table S4) using the primer set, and ran for 25 cycles (Table S5). For consensus ORs, template DNA was the hydrated synthesized OR, for native ORs, the template

DNA was the genomic DNA from chicken skeletal muscle (Zyagen #GC-314). Following amplification, we ran 1uL of product on a 1.5% agarose gel at 100V for 20 minutes, to confirm the amplified product. We then performed a purifying step and added 200uL of PB buffer (Qiagen) directly to the PCR strip tube. We then mixed the PB buffer and the PCR product and transferred to a Qiagen MinElute column and collection tube (Qiagen), followed by a 30 second spin at 15,000 RPM, and discarded the flow through. We then added 750uL of PE buffer followed by a 30 second spin, and then a 2 minute dry spin, discarding the flow through each time. We then added 10uL of elution buffer directly to the column and transferred the column to a 1.5mL microcentrifuge tube. We then spun the column down for 1 minute. For restriction enzyme digestion, we used 100ng/uL concentration of vector DNA, and added reagents (Table S6), including MluI and NotI high fidelity enzymes (New England Biolabs), and digested for 20 minutes at 37C. Following restriction enzyme digestion, we then purified the digested product. We added 200uL PB buffer to the PCR strip tubes, mixed thoroughly, and then transferred the mixture to the same MinElute column as before. We then spun for 30 seconds at 15,000 RPM and discarded the flow-through. We then added 750uL or 3.66M guanidine hydrochloride aqueous solution, made from 35g of guanidine hydrochloride in 100mL Gibco water. We then spun for 30 seconds and discarded the flow-through. We then added 750uL PE buffer (Qiagen) and spun for 30 seconds. Then, a second round of 750uL PE buffer, spinning for 30 seconds, followed by a 2 minute dry spin, discarding flow-through with each spin. We then transferred the MinElute spin column to a 1.5mL microcentrifuge tube, and added 10uL of elution buffer directly to the column, and spun for 1 minute. We then used the purified digest to set up a ligation reaction (Table S7), and held tubes at room temperature for over 1 hour. We ligated inserts into the rhodopsin tagged pCI vector or a modified Lucy-tagged, FLAG-tagged, and rhodopsin-tagged pCI vector, which contains MluI and NotI restriction enzyme sites, a Rho tag, as well as ampicillin and carbencillin resistance.

We then transformed *E. coli* cells with the plasmid with the ligated insert. We first thawed 20uL of competent cells and added 2.5uL of the ligated product. We held the incubating cells on ice for over 10 minutes, and then pipetted the cells onto an LB-ampicillin or LB-carbenicillin plate. We then let the colonies grow for 37 C overnight. To check for successful insertion of DNA into plasmid and transformation, we picked colonies and soaked in 20uL of Gibco water. We then performed colony PCR using primers designed for pCI (pCI 5'A: CTCCACAGGTGTCCACTC, pCI 3'A: CACTGCATTCTAGTTGTGG, Table S8) and ran for 25 cycles (Table S9). We then ran 1uL of PCR product with 5uL 1.2X loading dye (New England Biolabs) and checked for a band of the appropriate size on a 1% gel. We then added 10uL of *E. coli* with the appropriate insert to 4.5mL 2XYT-ampicillin or 2XYT- carbenicillin (100ug/mL) in a 17x100mm culture tube and shook bacteria for 18 hours at 37C. We then stored remaining volume from the colonies with the correct insert in 15-40% glycerol at -80C. Following incubation, we spun down the liquid culture for 30 seconds at 15,000 RPM twice in a 2mL tube and discarded the supernatant, retaining only the bacteria pellet. We then miniprepped plasmids with the ZymoPURE Plasmid Miniprep kit (Cat. # D4212, Zymogen, USA).

***Addition of Givaudan sequence to OR***

One challenge with studying native ORs are their low expression in heterologous cells. Comparisons in odor ligand selectivity and sensitivity between native ORs and consensus ORs are made difficult by low native OR expression that led to low native OR response. Previous studies have utilized the addition or modification of N-terminal and C-terminal tags to improve

OR trafficking from the ER to the cell surface (*82*). Studies have shown that basic amino acid residues, such as arginine and lysine placed at the C-terminus is beneficial for cell surface expression (*75*). Here, we engineered chicken native gamma-c OR receptors to investigate the potential improvement in the native OR receptors response.

We engineered ORs to modify their C-terminal amino acid sequences by replacing them with the basic Givaudan sequence RNKEVKKAIKRLFKRKCCRRR. To achieve this, we first obtained the nucleotide sequences of native ORs through DNA sequencing. Next, we designed primers to bridge the C-terminal region of the native ORs with the Givaudan sequence (*32*). Native ORs featured a conserved landmark sequence, NPXIYXXRN, on their C-terminus. The primer was specifically designed as a reverse complement to the junction between the native OR sequence and the Givaudan sequence. In this primer, the 3′ end included the reverse complement of the nucleotide sequence corresponding to the NPXIYXXRN region of the native OR. These nucleotides were selected to achieve a denaturation temperature of 56°C or 58°C, with adenine (A) and thymine (T) contributing 2°C each and cytosine (C) and guanine (G) contributing 4°C each. Preceding these nucleotides, the primer included the sequence GATCGCTTTTTCACTTCTTT, which encodes the reverse complement of the Givaudan amino acid sequence beyond the overlapping RN landmark. After obtaining OR specific bridging primers, we proceeded with OR cloning to achieve purified plasmid (previously described in STAR methods).

***In vitro cell culture: preparation of stock Hana3A cell line***

We kept Hana3A cell stocks frozen in -80 C freezers and followed a previously described protocol for cell preparation (*76*). We thawed frozen cells in 1mL tubes in a 37 C water bath.

After thawing, we transferred the entire stock to 6mL Minimum Essential Medium Eagle (MEM) (Cat # 10-010CV, Corning, Mediatech. Inc., VA, USA) supplemented with 10% Fetal Bovine Serum (FBS) (M10 solution) in a 15mL conical tube. We then centrifuged at 1000 rpm at room temperature and collected Hana3A cells in a pellet. We then aspirated the M10, leaving the pellet, and resuspended cells with 10mL M10 with and 0.5% penicillin-streptomycin (Gibco) and 0.5% amphotericin B (Gibco) on a 100-mm cell culture plate. We then cultured cells in a 37 C incubator with a water bath and 5% CO2 overnight. The next day, we checked cells with a phase contrast microscope to check for cell health. Ideal health has cells spread out uniformly across the surface of the culture dish and with no signs of contamination. We then replace medium with fresh M10 with penicillin-streptomycin and amphotericin medium to maintain cell health.

Over the next few days, we checked cell density (Hana3A cell line divides roughly once per 24 hours) and cell health periodically using a phase contrast microscope. We waited until cells reached the desired confluency and then we were ready to passage cells to 96 well plates. First, we tilted the dish to aspirate all media from the cell culture dish, and then we pipette 10mL PBS onto cells to wash. We then aspirated the PBS from the edge of the dish, and then added 3mL 0.05% trypsin-EDTA to the cells to detach them from the bottom of the dish. To assist in detaching, we gently shake the dish to detach cells from the bottom. This gentle shaking lasts around 2 minutes, and can be observed both by eye and under a phase contrast microscope. Immediately after detaching, we neutralized the trypsin-EDTA by the addition of 5mL M10 to the dish. We then dissociate the cells from forming clumps by pipetting all 8mL up and down onto the plate. We then transfer the cells and media to a 15mL conical tube, and centrifuge for 5 minutes at 200g at room temperature. Following spin down, we aspirated all media (M10 and trypsin-EDTA), without disturbing the cell pellet. We then resuspend cells in 1mL M10 to both seed 96-well plates and for the maintenance of the cell line.

For cell line maintenance, we transfer 10mL M10 with penicillin-streptomycin and amphotericin to a new culture plate. We then transfer the desired amount of cells to continue the cell population onto the plate, and place in a 37C incubator with 5% CO2 for future experiments. For cells to be used for transfection, we transferred to 96-well plates. The amount of cell transferred to 96 well plates should remain consistent, with roughly 10% of cells on a 100% cell covered 35x10mm cell culture dish per each 96-well plate used. So, for example, if the surface of a culture plate was covered 100% in cells, and an assay used six 96-well plates, 600uL (60%) of the cells from the culture plate resuspended in 1mL M10 would be used for 96-well plate seeding.

We then added 5mL M10 (without penicillin-streptomycin and amphotericin) per 96-well plate (Corning) used to a reagent reservoir (for example, an experimental design with six 96-well plates would use 30mL M10) and added the appropriate quantity of cells to the reservoir as well. We additionally added 60uL poly-D-lysine to the reservoir per 96-well plate (for example, 360uL poly-D-lysine for a design with six 96-well plates). We then mixed all of cells, media, and reagents. Using a multichannel pipettor, we then pipetted 50uL of the mixture from the reservoir to the each well of the 96-well plate. We then cultured the cells in the 96-well plate (s) for 24 hours in the 37C incubator with 5% CO2.

***Transfection of OR plasmid***

Prior to transfection, we prepared all necessary plasmids through the ZymoPURE Plasmid Miniprep kit (Cat. # D4212, Zymogen, USA). Necessary plasmids include ORs of interest,

RTP1S to assist in membrane transport (*77*), pGloSensor-20F (Cat # E1171, Promega Co., WI, USA) to encode for the cAMP-responsive element, and empty pCI mammalian expression vector (Cat # E1731, Promega Co., WI, USA) as a negative control. Rho-tagged ORs of interests are located in the pCI vector. We then observed the cells to be transfected for normal shape distribution, and a confluency of 30-50%. We then prepare two mixtures for each 96-well plate, first we prepared a DNA transfection mixture of 500uL MEM, 10uL pGLO, and 5uL RTP1S. After this is distributed, ORs are added to a final concentration of 100ng/uL per 96-well plate. We then add the second mixture of 500uL MEM with 20uL Lipofectamine 2000 (Cat # 11668019, Thermo Fisher Scientific Inc., MA, USA). Once the plasmids and Lipofectamine mixture are added together, we then incubate at room temperature for 15 minutes. Then for each 96-well plate, we added 5mL M10 to the mixture.

With the mixture prepared, we then removed the 96-well plates from the incubator and gently tapped the plates upside-down on sterile paper towels so that M10 from the previous day is removed from wells. Using a multichannel pipettor, we then added 50uL of the transfection mixture to each well. We then incubated cells with transfected plasmids for 24 hours in the 37C incubator with 5% $CO_2$.

***Loading stimulation buffer***

First, we observed cells under the microscope to check for roughly 50%-80% confluence and a healthy appearance. We then added 10mM HEPES and 15mM $NaN_3$ to HBSS to later add to the stimulation buffer. For a 500mL bottle of HBSS, we added 5mL 1M HEPES and 5mL 1.5M $NaN_3$. We then removed GloSensor cAMP Reagent (glo green) (Cat # E1291, Promega, WI, USA) from -80C storage and warmed to room temperature. We added 75uL glo green reagent to

2.76mL HBSS to create a stimulation buffer. We then removed the transfection media mixture from the cells by gently inverting and tapping 96-well plates onto sterile paper towels in a safety cabinet. We then used a multichannel pipettor to distribute 25uL of the stimulation buffer to each well. We then placed the 96-well plate at room temperature covered in aluminum foil in a drawer for 2 hours. Glo green reagent is light sensitive and we kept plates in a dark, odorless environment.

During incubation, we prepared odorants. Initially, we diluted odorants to 100mM concentrations in 95% ethanol and stored at -20C. We then added odors to the stimulation buffer at the desired concentration for odor exposure to ORs. For example, we would serially dilute 100mM working solutions to concentrations of 300uM, 100uM, 30uM, 10uM, 3uM, 1uM, in continuum, at the time of stimulation. Each OR in each experiment also received a no odor negative control, with only stimulation medium, to assess background cAMP activity levels. For odorants with sulfur, we added 30uM $CuCl_2$ to the odorant stimulation buffer mixture.

Following incubation, we transferred each 96-well plate to the CLARIOstar Plus multi-mode plate reader (BMG LABTECH). We then take a baseline measurement of fluorescence activity of the cells prior to the addition of odorants. Following the blank measurement, using a multichannel pipettor, we distributed 25uL of the odorant and stimulation medium mixture to each well in the 96-well plate. Immediately following odorant exposure, we placed the 96-well plate back in the CLARIOstar plate reader and began reading cAMP activity for 15 minutes, at 10 cycles, or one cycle per 90 seconds.

***Cell-based assay data analysis***

We analyzed plate reader data using custom python scripts. We normalized fluorescence activity to the blank zero time point measurement prior to odor stimulation, and subtracted this value by 10 to allow for 0 to signify a lack of response. We analyzed dose-response curves to ORs by fitting a least squares function to the data in GraphPrism 10 Version 10.4.1 (532).

***OR cell surface expression***

Flow cytometry was performed to assess the cell-surface expression of ORs. HEK293T cells were plated in 6 well plate (Corning) at a density of approximately $3.5 \times 10^5$ cells (2.5% confluency per well) and cultured overnight. After 18–24 hours, OR plasmids (1,000 ng), which were N-terminally tagged with the first 20 amino acids of human rhodopsin (rho-tag) in the pCI mammalian expression vector (Promega), were transfected along with 200 ng of RTP1S and 10 ng of eGFP using Lipofectamine 2000 (Cat # 11668019, Thermo Fisher Scientific, MA, USA).

At 18–24 hours post-transfection, cells were detached using Cell Stripper (Cat # 25-056-CI, Corning, Mediatech Inc., VA, USA) and resuspended in ice-cold Phosphate Buffered Saline (PBS) (Cat # SH30256.01, Cytiva, HyClone Laboratories, UT, USA) supplemented with 15 mM Sodium Azide ($NaN_3$) (Cat # S2002, Sigma-Aldrich, MO, USA) and 2% Fetal Bovine Serum (FBS) (Cat # SH3008802HI, Cytiva, HyClone Laboratories, UT, USA). The cell suspension was transferred to 5 ml round-bottom polystyrene tubes (BD), centrifuged at 4°C, and resuspended again in PBS containing 15 mM $NaN_3$ and 2% FBS. Cells were then incubated with a primary antibody (1/400 dilution, mouse anti-rhodopsin clone 4D2, MABN15, Sigma-Aldrich, MO, USA) for 30 minutes, followed by washing with PBS containing 15 mM $NaN_3$ and 2% FBS.

After another centrifugation step, cells were stained with a secondary antibody (1/200 dilution, phycoerythrin-conjugated donkey anti-mouse F (ab′)2 fragment, 715-116-150, Jackson Immunologicals) for 30 minutes in the dark. To distinguish dead cells, 7-amino-actinomycin D (1/500 dilution, 129935, Calbiochem) was added. The samples were immediately analyzed using a BD FACSCanto II flow cytometer, with gating applied to select GFP-positive, single, spherical, and viable cells. Phycoerythrin fluorescence intensities were quantified and visualized using FlowJo v10.8.1. An empty pCI plasmid served as the negative control.

***Live chicken exposure to pyrazines***

We collected 64 week old white leghorn chickens and placed in boxes with airflow through an external vent (IACUC protocols 23-429 and 22-280). We allowed chickens to be in the box for one hour to acclimate the chickens from any outside odors they may have encountered prior to the experiment. Following one hour acclimation, we placed odors on blotting paper that was then placed inside a Sakura Tissue-Tek Uni-cassette system. Odors added to the cassette were either 1% acetophenone, 1% 2-isobutyl-3-methoxypyrazine, or 10% 2-isobutyl-3-methoxypyrazine, diluted in ethanol. No odor control chickens received blotting paper and cassette with no odor added. We then placed cassettes on the floor of the boxes with the chickens. Chickens were standing on a grated floor at the bottom of the box, and we placed cassettes below the grated floor to prevent chickens from interfering with cassettes.

Following one hour of odor exposure, we then sacrificed chickens. We then dissected chickens were to obtain posterior olfactory epithelium tissue. We embedded posterior olfactory epithelium in OCT medium, and flash froze in liquid nitrogen prior to transfer to -80C. We then cut 18uM sections of the posterior olfactory epithelium using a Leica CM 1850 Cryostat and placed on VWR superfrost slides, and stored at -80C.

***Odorant receptor structure generation***

Native chicken gamma-c OR structure was generated using AlphaFold3 methods published previously (*86*). We compiled odorant receptor protein sequence together with $G_{olf}$ peptide and odorant SMILE into standardized JSON configuration files. Twenty independent structural prediction of the OR-$G_{olf}$-ligand complex were generated using AlphaFold3, and we selected the structural prediction with highest ranking score (most confidence) to visualize. For visualization, we utilized UCSF ChimeraX (*87*) to remove $G_{olf}$ and color code receptor based on their percent identity.

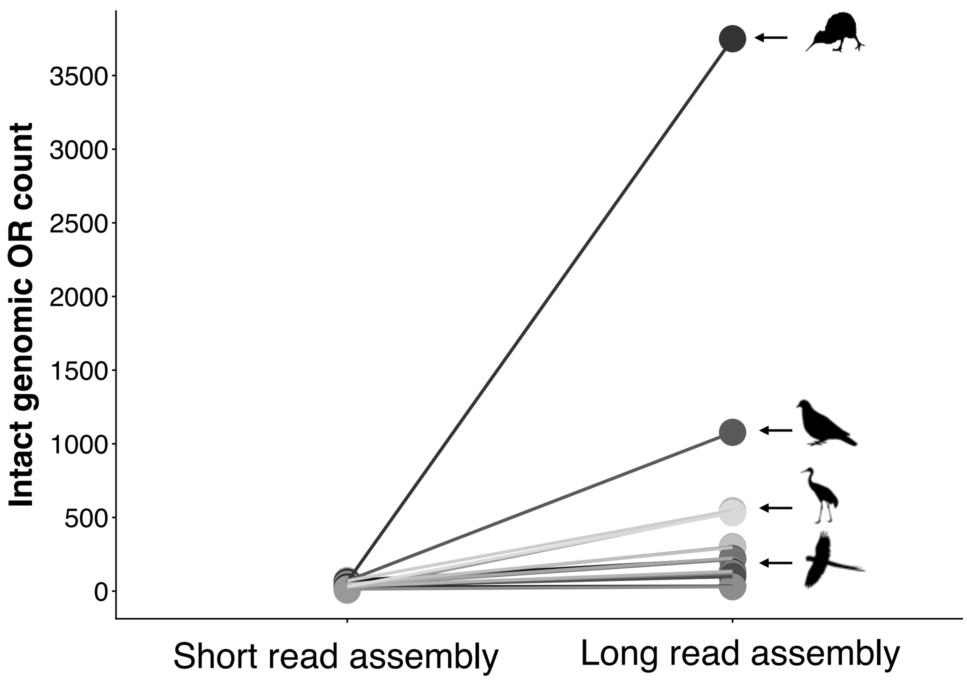


**Fig. S1.**
**Long read assemblies greatly increase the genomic OR repertoire of birds.**
Each circle represents the number of ORs from the same species found in short and long read assemblies. Short read assembly values are from Policarpo et al.[12]

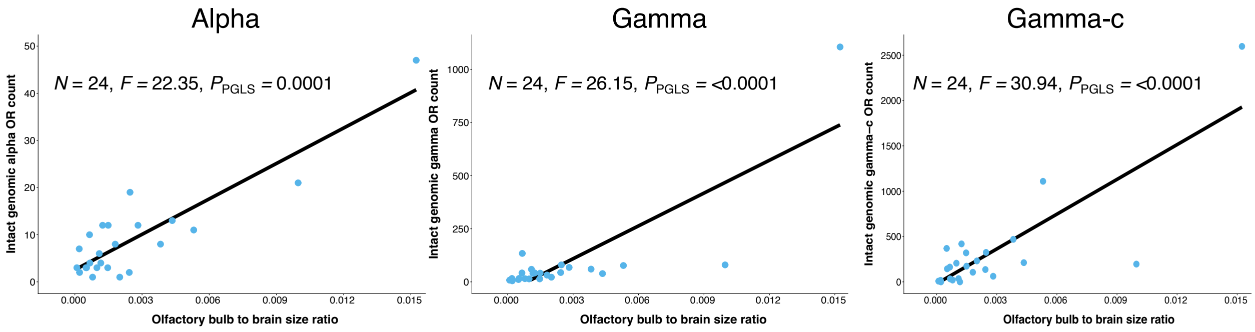


**Fig. S2.**

**Olfactory bulb size is correlated with OR counts of OR subfamilies.** This relationship suggests that each individual subfamily may be involved in smell, including the gamma-c ORs. Correlation measured with phylogenetic generalized least squares test.

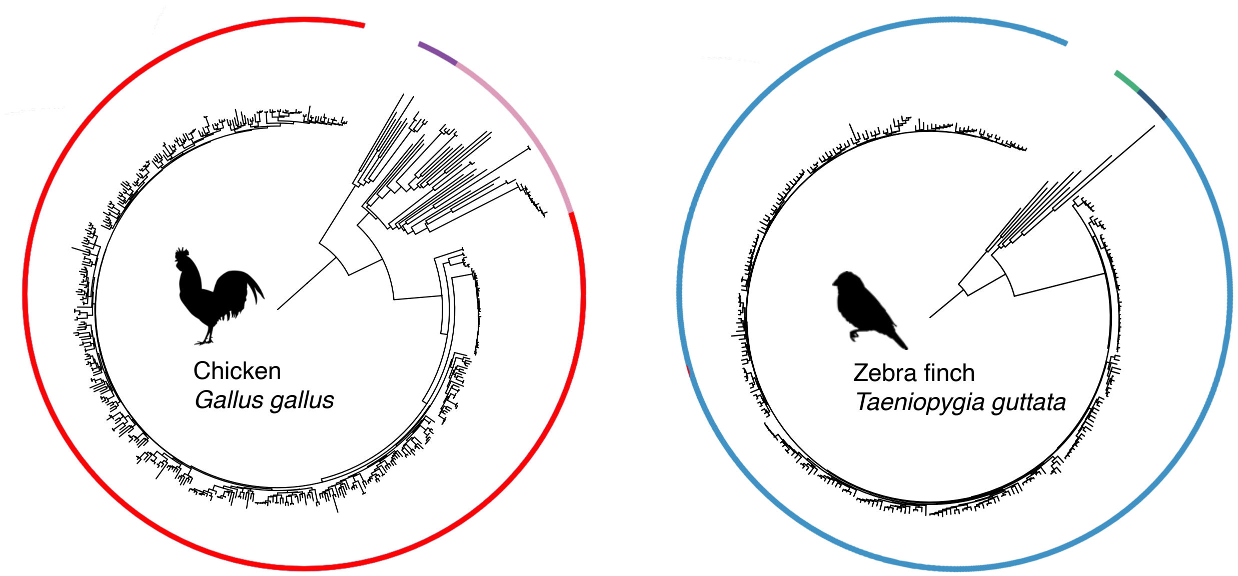


**Fig. S3.**
**Genomic OR repertoires from model bird species.** Phylogeny of 473 intact genomic ORs in the chicken (right). Purple shows alpha receptors, pink shows gamma receptors, red shows gamma-c ORs. Phylogeny of 278 zebra finch ORs (left). Green indicates alpha ORs, dark blue gamma, and light blue gamma-c (class II) ORs. Bird (black) and mammal (orange) silhouettes from Phylopic.

\

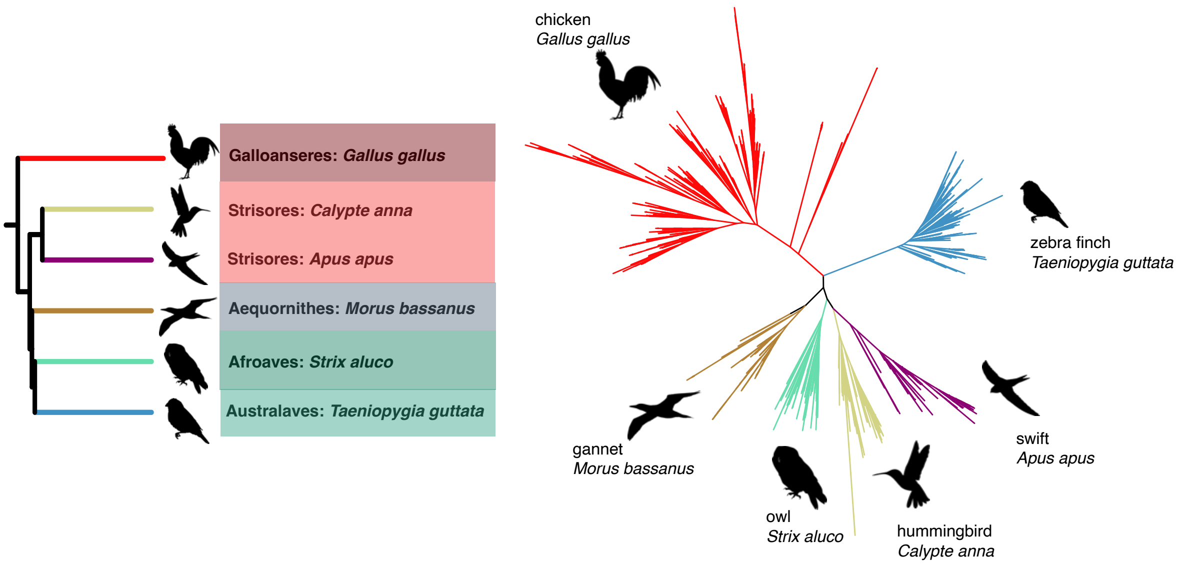


**Fig. S4.**
**Gamma-c ORs form family-specific clades.** Phylogeny of six bird species representing six families and five orders (left). Phylogeny of gamma-c OR genomic repertoires from the six species (right) form species-specific clades

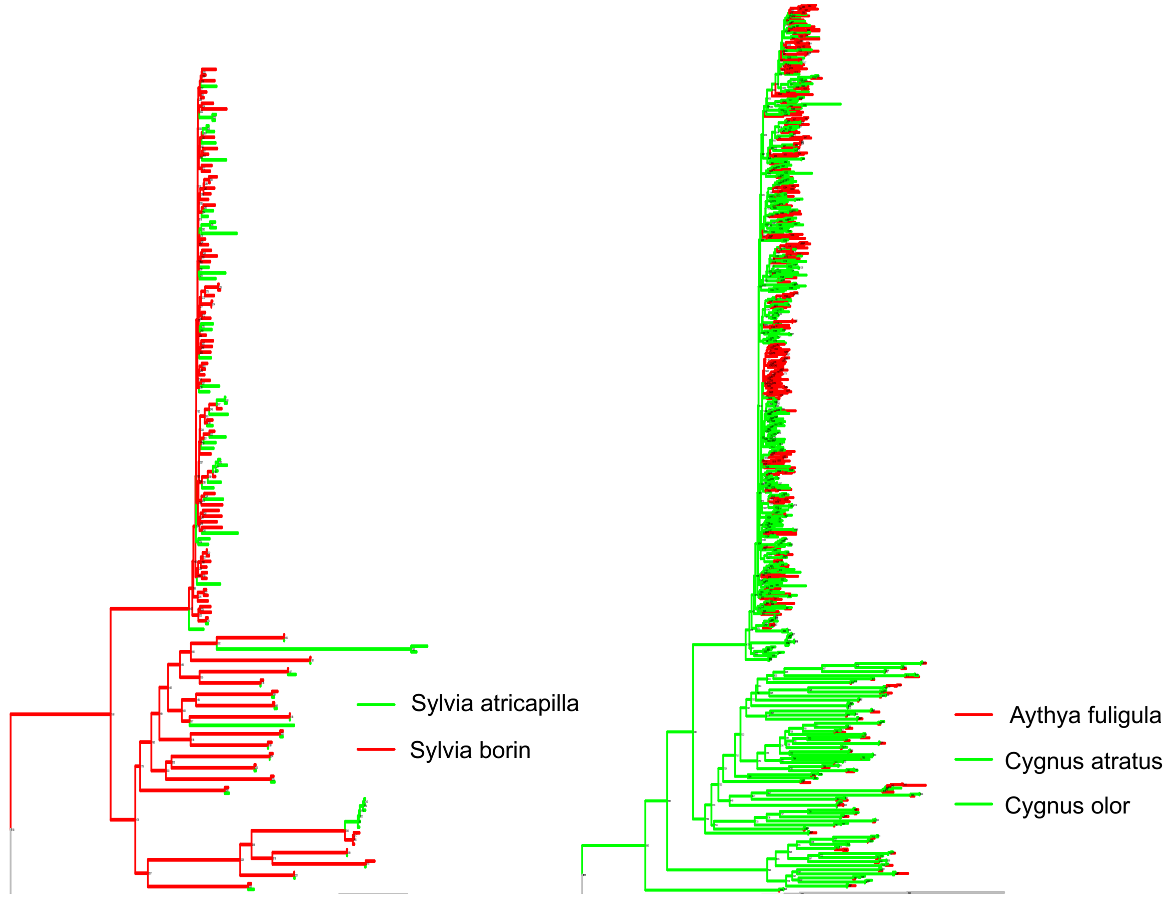


**Fig. S5.**
**Closely related bird species within the same bird family do not show monophyletic gamma-c OR clades.** Two species in the Old World warbler family (Sylviidae) as well as tufted duck (*Aythya fuligula*) and two swan species (*Cygnus atratus* and *C. olor*) show interdigitation of gamma-c OR in the phylogeny.

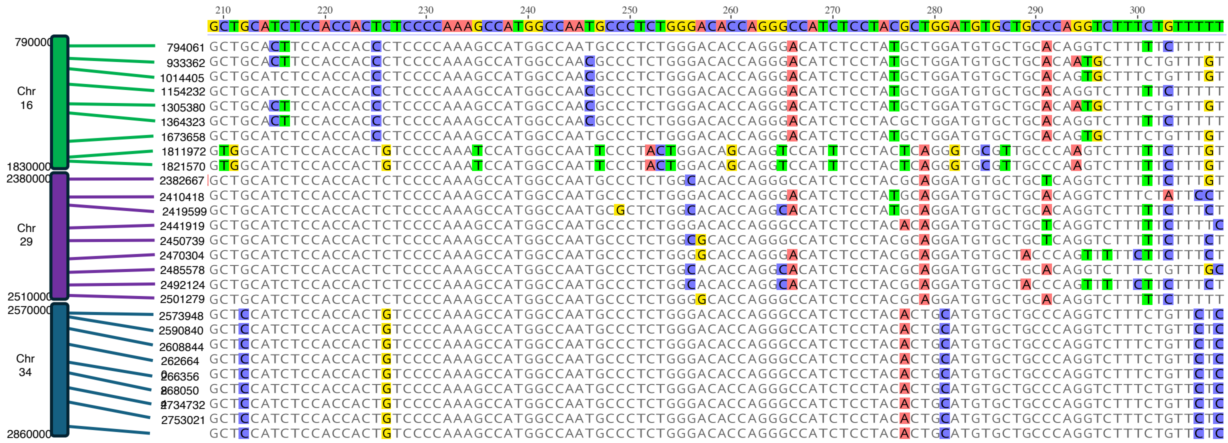

**Fig. S6.**

**Chromosome-specific signatures of gamma-c ORs in chicken.** Coordinates of representative chicken gamma-c ORs from three dot chromosomes are shown. Nucleotides divergent from consensus sequence are colored. Substitutions largely follow chromosome-specific patterns.

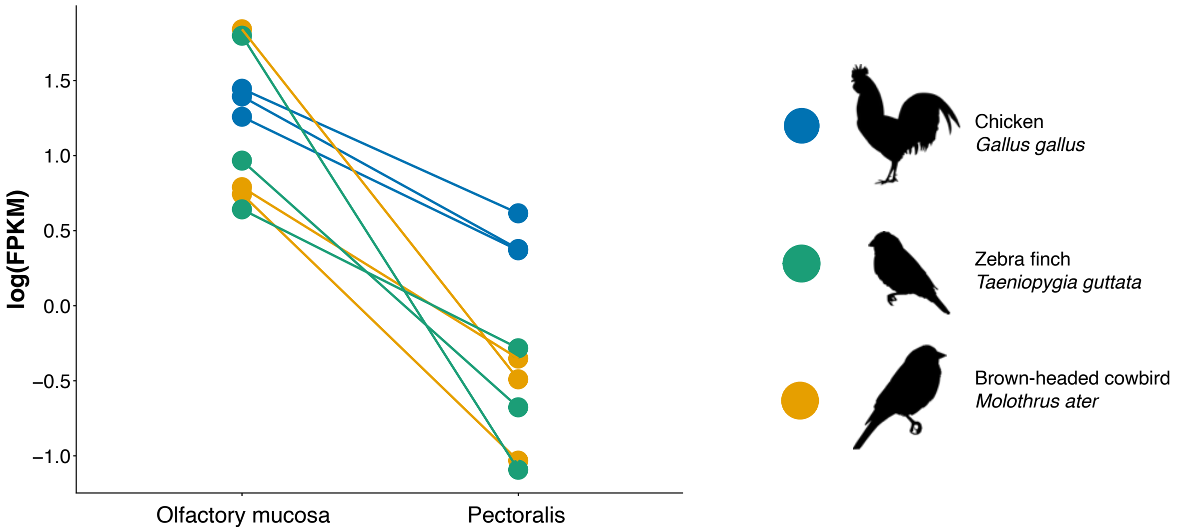


**Fig. S7.**
**Expression of OSN markers is higher in olfactory epithelium than pectoralis muscle.** We summed expression of three OSN positive markers, Cnga2, Omp, and Adcy3, and observed high levels in the olfactory mucosa samples. This suggests that we have isolated tissue containing OSNs. Additionally, samples with larger OSN marker expression consistently had larger OR expression (Fig. 3a).

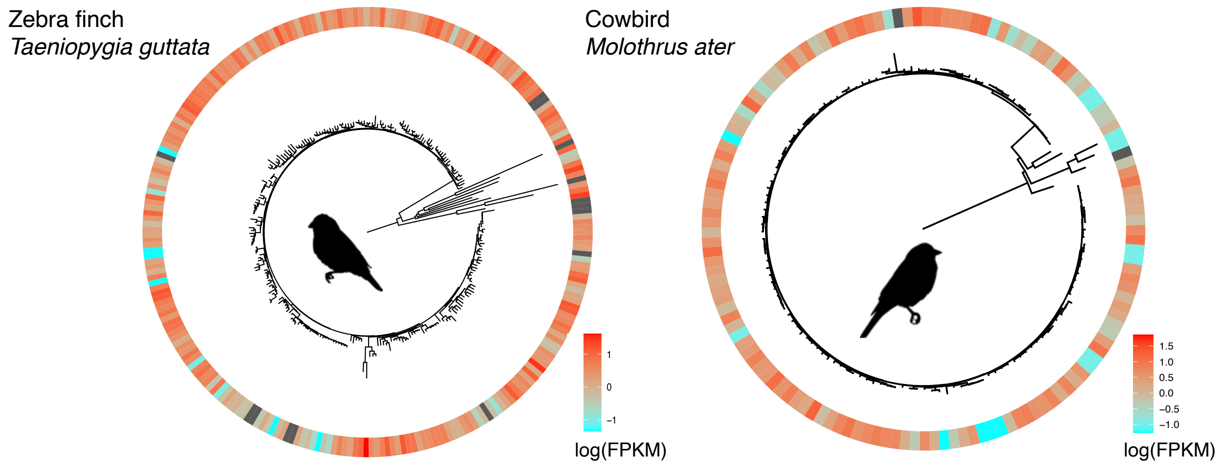


**Fig. S8.**

**OR expression levels of individual ORs in olfactory mucosa and pectoralis muscle tissue.** Zebra finch (*Taeniopygia guttata*, left) and cowbird (*Molothrus ater,* right) are depicted with a phylogeny of its intact genomic OR repertoire. The ring shows the olfactory mucosa expression. Cyan indicates low level of expression, red high level of expression.

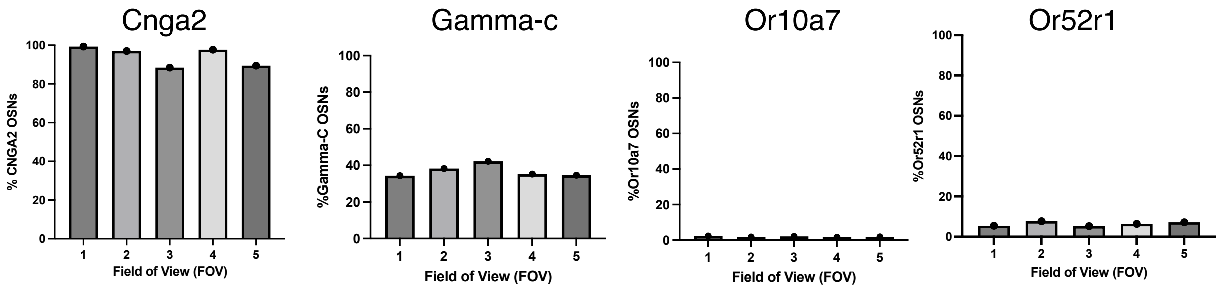


**Fig. S9.**

**Semiquantitative analysis of *in situ* hybridization signals for OSN marker Cnga2 and chicken ORs in the olfactory epithelium.** Five randomly selected fields of view were analyzed for each probe staining to quantify the percentage of olfactory sensory neurons expressing the canonical OSN marker Cnga2, and different chicken ORs, gamma-c, Or10a7, and Or52r1 within the chicken olfactory epithelium.

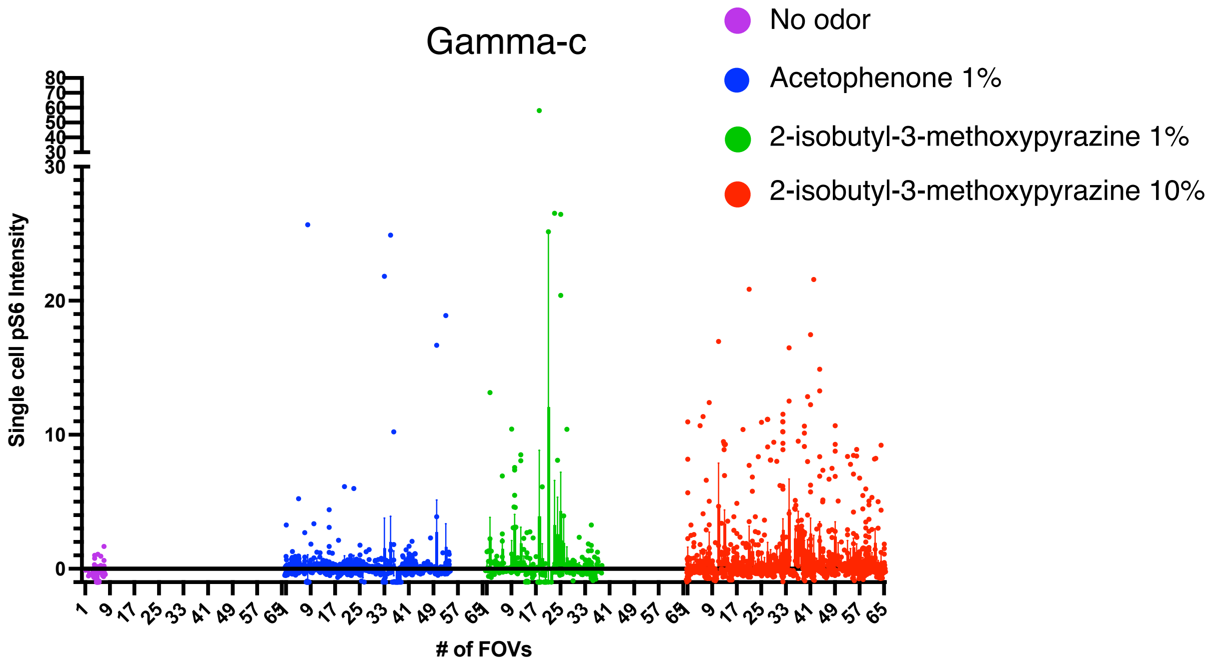


**Fig. S10.**
**pS6 intensity semi-quantitative measurement in individual gamma-c–positive neurons.** Each dot represents the pS6 fluorescence intensity measured per cell within the identified gamma-c–positive population. Semi-quantification was performed at the single-cell level, and data are expressed as relative fluorescence intensity normalized to background signal.

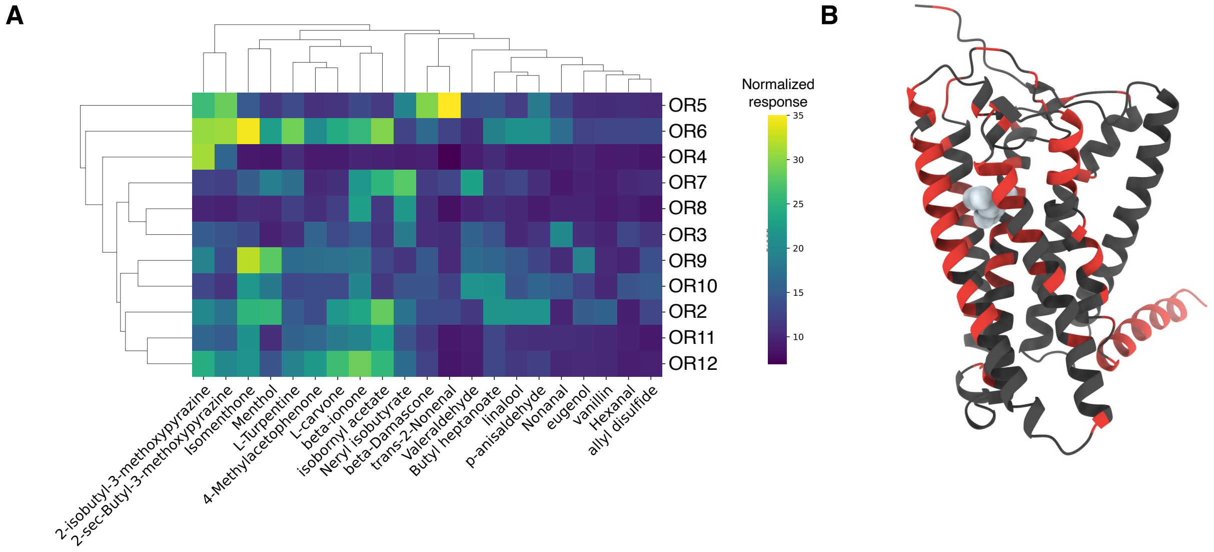


**Fig. S11. Chicken gamma-c ORs respond to a variety of odors, and diverse nucleotide regions surround the predicted odor binding pocket. (A)** Heatmap of odor responses to 11 chicken gamma-c ORs. ORs 2, 3, and 4 are the same as shown in Fig. 5B. Assays conducted in duplicate and normalized by the fluorescence at the first time point. (B) AlphaFold3 structure of chicken gamma-c OR is bound to $G_{olf}$ (not shown) in an active-like configuration. Homogenized region is colored with black and diverse region is colored with red.

| Reagent | Volume (uL) |
|---|---|
| 10X Buffer | 1 |
| 2 mM dNTPs | 1 |
| 5 uM forward primer | 1 |
| 5 uM reverse primer | 1 |
| Taq DNA polymerase (Qiagen) | 0.05 |
| Distilled water | 6 |
| Plasmid DNA (1 ng/uL) in elution buffer | 1 |
| Total | 11.05 |

**Table S1.**
PCR reaction mix to amplify probe template DNA.

| Step | Temperature | Time |
|---|---|---|
| Initial denaturation | 95°C | 15 minutes |
| 25 cycles | 95°C | 15 seconds |
| | 55°C | 15 seconds |
| | 72°C | 1 minute (per template DNA kilobase) |
| Final extension | 72°C | 5 minutes |
| Hold | 10°C | |

**Table S2.**
PCR cycling conditions to amplify probe template DNA.

| **Reagent** | **Volume (uL)** |
|---|---|
| Purified template DNA | 1 uL |
| 5x RNA polymerase buffer | 2 uL |
| DIG RNA labeling mix | 1 uL |
| T3 RNA polymerase | 0.5 uL |
| RNase inhibitor mix | 0.5 uL |
| DTT (100 mM) | 1 uL |
| Nuclease-free water | 4 uL |
| Total | 10 uL |

**Table S3.**
Transcription reaction mix to synthesize RNA probe.

| Reagent | Volume (uL) |
|---|---|
| 5X Phusion buffer | 2 |
| 2 mM dNTPs | 1 |
| 5 uM forward primer | 1 |
| 5 uM reverse primer | 1 |
| Phusion DNA polymerase | 0.1 |
| Distilled water | 5 |
| Plasmid DNA (1 ng/uL) in elution buffer | 1 |
| Total | 11.1 |

**Table S4.**
PCR reaction mix to amplify ORs from genomic DNA.

| Step | Temperature | Time |
|---|---|---|
| Initial denaturation | 98°C | 30 seconds |
| 30 cycles | 98°C | 5 seconds |
| | 55-65°C (delta T = -0.5°C) | 15 seconds |
| | 72°C | 1 minute (per template DNA kilobase) |
| 15 cycles | 98°C | 5 seconds |
| | 55°C | 15 seconds |
| | 72°C | 1 minute (per template DNA kilobase) |
| Final extension | 72°C | 5 minutes |
| Hold | 10°C | |

**Table S5.**
PCR cycling conditions to amplify OR from genomic DNA.

| Reagent | Volume (uL) |
|---|---|
| DNA (100 ng/uL) | 9 |
| Cutsmart buffer | 2 |
| MluI-HF | 0.5 |
| NotI-HF | 0.5 |
| Distilled water | 8 |
| Total | 20 |

**Table S6.**

Restriction enzyme digest reaction mix to cut vector.

| Reagent | Volume (uL) |
|---|---|
| DNA insert | 2 |
| Vector (Rho-pCI) | 0.5 |
| T4 ligase | 0.5 |
| 10X buffer | 0.5 |
| Distilled water | 1.5 |
| Total | 5 |

**Table S7.**
Ligation reaction mix to insert DNA segment.

| Reagent | Volume (uL) |
|---|---|
| 10X buffer | 1 |
| 2 mM dNTPs | 1 |
| 5 uM forward primer | 1 |
| 5 uM reverse primer | 1 |
| Taq DNA polymerase | 0.1 |
| Distilled water | 5 |
| Bacteria colony in distilled water | 2 |
| Total | 11.1 |

**Table S8.**
Colony PCR reaction mix to confirm bacterial transformation.

| Step | Temperature | Time |
|---|---|---|
| Initial denaturation | 95°C | 15 minutes |
| 25 cycles | 95°C | 15 seconds |
| | 55°C (delta T = -0.5°C) | 15 seconds |
| | 72°C | 1 minute (per template DNA kilobase) |
| Final extension | 72°C | 5 minutes |
| Hold | 10°C | |

**Table S9.**
Colony PCR cycling conditions to confirm bacterial transformation.

**Data S1. (separate file)**
Results of running GENECONV tool on chicken (GCA_024206055.2) gamma-c OR repertoire.

**Data S2. (separate file)**
List of accession numbers of genomes with reported OR repertoires.

**Data S3. (separate file)**
Python script for filtering OR pseudogenes.

**Data S4. (separate file)**
Primers used for generating probes used in *in situ* hybridization.

**Data S5. (separate file)**
Primers used for cloning native chicken olfactory receptors.

**Data S6. (separate file)**
Alignment of gamma-c ORs for species in Fig. 2A.

**Data S7. (separate file)**
Bird OR counts and data used for Fig. 1A, B.

**Data S8. (separate file)**
Human OR sequence used for transmembrane domain identification.

**Data S9. (separate file)**
Outgroup sequences used for OR phylogenetic relationships.

**Data S10. (separate file)**
BLAST query file used for OR genome search.

**Data S11. (separate file)**
Perl script for OR BLAST search of genome.

**Data S12. (separate file)**
Perl script for locating OR open reading frame.

**Data S13. (separate file)**
Perl script locating OR open reading frame with print out.

**Data S14. (separate file)**
Perl script for obtaining OR open reading frame coordinates.

**Data S15. (separate file)**
Perl script for finding correct start Methionine.

**Data S16. (separate file)**
Perl script for writing out final open reading frame coordinates.

**Data S17. (separate file)**
Sequences of all chicken (GCA_024206055.2) gamma-c ORs.

**Data S18. (separate file)**
Phylogeny of all bird species in Fig. 1A.

**Data S19. (separate file)**
Unix commands for running OR genome search.

**Data S20. (separate file)**
R code for filtering pseudogenes from OR genome search by length and genome overlap.

**Data S21. (separate file)**
Chicken OR consensus sequence and native ORs used in Fig. 4B, 4D.

**Data S22. (separate file)**
Human OR sequences used in Fig. 2B.

**Data S23. (separate file)**
Sequence used for Alphafold3 structure.

**Data S24. (separate file)**
Chicken gamma-c OR sequences used in fig. S11A.

**Data S25. (separate file)**
Sequence of chicken OR51E2 sequence.
