## Supplementary material for "Unique evolutionary radiation of odorant receptors in birds": DataS3_filter_pseudogene.py

```
import numpy as np 
import pandas as pd 
import os
import argparse

from filter_pseudogene_class import FilterPseudoGeneParams, FilterPseudoGene


parser = argparse.ArgumentParser()
parser.add_argument('--save_data', 
                    help = "boolean to save data", 
                    default=False)
parser.add_argument('--save_fasta', 
                    help = "boolean to save data", 
                    default=True)
parser.add_argument('--output_path', 
                    help = "file path to output data", 
                    default='./')
parser.add_argument('--query_file_path', 
                    help = "file path of .fasta to be queried ", 
                    required=True)
parser.add_argument('--domain_file_path', 
                    help = "file path of domain info", 
                    default = 'HumanORTMD.txt')
args = parser.parse_args()


# instantiate parameters
filter_params = FilterPseudoGeneParams()   
filter_params.save_data = args.save_data 
filter_params.save_fasta = args.save_fasta
filter_params.output_path = args.output_path
filter_params.query_file = args.query_file_path
filter_params.domain_file = args.domain_file_path


# instantiate class and run filter_genes 
filter_OR = FilterPseudoGene(filter_params)
filter_OR.filter_genes()
```
