## Supplementary material for "Unique evolutionary radiation of odorant receptors in birds": DataS11_Step_1_ab_tblastN_ORs.pl

```
#!/usr/bin/perl -w
use strict;
use warnings;    


use Bio::SearchIO;
use Bio::Seq;
use Bio::Tools::Run::StandAloneBlast;


my $data="chickendb"; # <ferret genome> 
my $query="OR_query_mini_ORN_gharial.fasta";
my $e="1e-20";
my $out=$query;   $out=~s/\.fasta/\.out/g; # have the output be the same as the query name but replace .fasta with .out
my $pro="tblastn";


system("tblastn -evalue $e -query $query -db $data -out $out -outfmt 5 -num_alignments 200000 -num_descriptions 200000 -num_threads 20");


my $input=$out;
my $output=$input;
$output=~s/\.out/\.sum/;


  my $file = $input;
  my $in = new Bio::SearchIO(-format =>'blastxml',
                             -file =>$file);
  my $num = $in->result_count;  


  open (OUT, ">>$output");

  print OUT "Query\/Query_length\/Hit\/Hit_length\/E-value\/Bit score\/Percent_identity\/Number_indentity\/Query_Start\/Query_End\/Hit_Start\/Hit_END\/Query_strand\/Hit_strand\n";

     while( my $r = $in->next_result )
   { 
     
     while( my $h = $r->next_hit ) 
     { 
   
         while( my $hsp = $h->next_hsp ) 
                 {
                      my $queryname=$r->query_name;
                       $queryname=~s/\|/\//g;
                       print OUT $queryname,";", " ", $r->query_description,"\/"," ",$r->query_length,"\/";

                       print OUT $h->name, "\/"; 
                       print OUT $hsp->length('total'),"\/", " ",  $hsp->evalue,"\/", $hsp->score, "\/",$hsp->percent_identity ,"\/",$hsp->num_identical,"\/",$hsp->query->start,"\/", " ", $hsp->query->end,"\/"," ", $hsp->hit->start,"\/", " ", $hsp->hit->end,"\/", " ", $hsp->query->strand,"\/", " ",$hsp->hit->strand,"\n";

                 }
                           

      }
}
```
