## Supplementary material for "Unique evolutionary radiation of odorant receptors in birds": DataS12_step_3_a_Find_ORF-ABmod.pl

```
#!/usr/local/bin/perl
#!/usr/bin/perl -w
# GeneFinder program

my $faspath="step_2_results.olfacUniqueBlastHits.stranded.fasta";
# needs to be in the directory, not absolute path from another place
# these are unique (step1c),stranded(revcomp as appropriate) and have been extended 300bp (100 codons) in each direction

my $expect_length=810;                                # length of the expected gene sequences

my $Output_file_name=$faspath;
   $Output_file_name='ORF_longThan_'.$expect_length.'_bp_'.$faspath; 

open (FILE,"$faspath");

open (OUT, ">>NO_ambu.fas");


system ("cls");


       
	
        while( <FILE> )
	{
		     my $line=$_;  chomp $line;     
                 if  ($line=~/^>/)   {print OUT "$line\n";next;}
                 else 
                 {
                 $line=~s/-//g;
                  $line=~s/R|Y|K|M|S|W/N/g;    
                   print OUT "$line\n";
                 }
	 }  
	close FILE;
     

my $input_file_name='./NO_ambu.fas';

# this makes a file with no ambiguities (all are changed to N by the above code # 


open(FILE,"$input_file_name");
open (OUT, ">>$Output_file_name");


     my $count_number; 
     my $species;
     my %sequences;
	
        while( <FILE> )
	{
		    my $line=$_;  
                if( $line =~ /^(>.+\n)/ )
                {
                        $species = $1;
                        $sequences{$species} = $species;   
                }
                else
                   {
                            $sequences{$species} .= $line;                      
                   }

	 }
	close FILE;


   foreach my $spe (keys %sequences)

{ 
   
 my $DNA_seq=$sequences{$spe};

my $sequenceEntry =$sequences{$spe};

my $sequenceTitle = "";
if ($sequenceEntry =~ m/(>[^\n]+)/){
   $sequenceTitle = $1;
   $sequenceTitle =~ s/>//;
}
else {
    die( "A FASTA sequence title was not found." );
}


$sequenceEntry =~ s/>[^\n]+//;

$sequenceEntry =~ tr/GATCN/gatcn/;

$sequenceEntry =~ s/[^gatcn]//g;

my @arrayOfORFs = ();

my @startsRF1 =();
my @startsRF2 =();
my @startsRF3 =();
my @stopsRF1 = ();
my @stopsRF2 = ();
my @stopsRF3 = ();

while ($sequenceEntry =~ m/atg/gi){
    my $matchPosition = pos($sequenceEntry) - 3;

    if (($matchPosition % 3) == 0) {
	push (@startsRF1, $matchPosition);
    }

    elsif ((($matchPosition + 2) % 3) == 0) {
	push (@startsRF2, $matchPosition);
    }
    
    else {
	push (@startsRF3, $matchPosition);
    }
}

while ($sequenceEntry =~ m/tag|taa|tga/gi){
    my $matchPosition = pos($sequenceEntry);
    if (($matchPosition % 3) == 0) {
	push (@stopsRF1, $matchPosition);
    }
    elsif ((($matchPosition + 2) % 3) == 0) {
	push (@stopsRF2, $matchPosition);
    }
    else {
	push (@stopsRF3, $matchPosition);
    }
}

my $codonRange = "";
my $startPosition = 0;
my $stopPosition = 0;

@startsRF1 = reverse(@startsRF1);
@stopsRF1 = reverse(@stopsRF1);

while (scalar(@startsRF1) > 0) {
    $codonRange = "";
    
    $startPosition = pop(@startsRF1);

    if ($startPosition < $stopPosition) {
	next;
    }
  
    while (scalar(@stopsRF1) > 0) {
	$stopPosition = pop(@stopsRF1);
	if ($stopPosition > $startPosition) {
	    last;
	}
    }

    if ($stopPosition <= $startPosition) {

	$stopPosition = length($sequenceEntry) - (length($sequenceEntry) % 3);

	$codonRange = "+1 " . $startPosition . ".." . $stopPosition;
	push (@arrayOfORFs, $codonRange);
	last;
    }

    else {

	$codonRange = "+1 " . $startPosition . ".." . $stopPosition;
	push (@arrayOfORFs, $codonRange);
    }
}

$stopPosition = 0;
@startsRF2 = reverse(@startsRF2);
@stopsRF2 = reverse(@stopsRF2);
while (scalar(@startsRF2) > 0) {
    $codonRange = "";
    $startPosition = pop(@startsRF2);
    if ($startPosition < $stopPosition) {
	next;
    }
    while (scalar(@stopsRF2) > 0) {
	$stopPosition = pop(@stopsRF2);
	if ($stopPosition > $startPosition) {
	    last;
	}
    }
    if ($stopPosition <= $startPosition) {

	$stopPosition = length($sequenceEntry) - ((length($sequenceEntry) + 2) % 3);
	$codonRange = "+2 " . $startPosition . ".." . $stopPosition;
	push (@arrayOfORFs, $codonRange);
	last;
    }
    else {
	$codonRange = "+2 " . $startPosition . ".." . $stopPosition;
	push (@arrayOfORFs, $codonRange);
    }
}

$stopPosition = 0;
@startsRF3 = reverse(@startsRF3);
@stopsRF3 = reverse(@stopsRF3);
while (scalar(@startsRF3) > 0) {
    $codonRange = "";
    $startPosition = pop(@startsRF3);
    if ($startPosition < $stopPosition) {
	next;
    }
    while (scalar(@stopsRF3) > 0) {
	$stopPosition = pop(@stopsRF3);
	if ($stopPosition > $startPosition) {
	    last;
	}
    }
    if ($stopPosition <= $startPosition) {

	$stopPosition = length($sequenceEntry) - ((length($sequenceEntry) + 1) % 3);
	$codonRange = "+3 " . $startPosition . ".." . $stopPosition;
	push (@arrayOfORFs, $codonRange);
	last;
    }
    else {
	$codonRange = "+3 " . $startPosition . ".." . $stopPosition;
	push (@arrayOfORFs, $codonRange);
    }
}

# print OUT "Results for \"$sequenceTitle\"\n";
# print OUT "Length = " . length($sequenceEntry) . " bp\n";

foreach(@arrayOfORFs) 
 {
    
   
    $_ =~ m/([\+\-]\d)\s(\d+)\.\.(\d+)/;

    my $frame_infor=$1;  my $begin=$2; my $end=$3;
    my $seg=substr($sequenceEntry,$begin,($end-$begin));  my $len_seq=length($seg);
    if ($len_seq  >=  $expect_length)                                      #!!!!
    {
     
   #  print OUT "==============================================================\n";
   #  print OUT "==============================================================\n";
    chomp $spe; 
	my $numberAA=($end - $begin)/3;
    
  #  print OUT "--Reading frame $frame_infor " . ($begin + 1) . " to " . $end;     
  #  print OUT ", Length: " . ($end - $begin+1)."\n";
  #  print OUT "--SEQUENCES:  $seg\n";
    my $AA=GetAA($seg);  
	
	if ($AA!~/X/) {
	$count_number++;
	 print OUT "$spe-".$numberAA."_aa\n";
	 print OUT "$AA\n";
	               }
   #  print OUT ">>>>>>>>>>>>>>>>>>>>>>>>>>>>>>>>>>>>>>>>>>>>>>>>>>>>>>>>>>>>>>>>>>\n";
   #  print OUT ">>>>>>>>>>>>>>>>>>>>>>>>>>>>>>>>>>>>>>>>>>>>>>>>>>>>>>>>>>>>>>>>>>\n\n";
   }

 }


}

#print OUT "$count_number gene found\n";

system ("del NO_ambu.fas");


sub GetAA

{
 my $trans=shift;
 my $proseq;
  for( my $i=0; $i<(length($trans)-2); $i+=3 )
                        {
                                $codon = substr( $trans,$i,3 );
                                if( $codon eq 'TAG' or $codon eq 'TAA' or $codon eq 'TGA' )
                                {
                                        last;
                                }
                                if( $codon =~ /N/ )
                                {
                                        $proseq .= 'X';
                                	$cdsseq .= 'NNN';
                                }else{
                                	$proseq .= codon2aa($codon);
                                	
                                }
                        }
                       
                        return ($proseq);
}

sub codon2aa
{
	my($codon) = @_;
	if ( $codon =~ /GC./i) { return 'A' }
	elsif ( $codon =~ /TG[TC]/i) { return 'C' }
	elsif ( $codon =~ /GA[TC]/i) { return 'D' }
	elsif ( $codon =~ /GA[AG]/i) { return 'E' }
	elsif ( $codon =~ /TT[TC]/i) { return 'F' }
	elsif ( $codon =~ /GG./i) { return 'G' }
	elsif ( $codon =~ /CA[TC]/i) { return 'H' }
	elsif ( $codon =~ /AT[TCA]/i) { return 'I' }
	elsif ( $codon =~ /AA[AG]/i) { return 'K' }
	elsif ( $codon =~ /TT[AG]|CT./i) { return 'L' }
	elsif ( $codon =~ /ATG/i) { return 'M' }
	elsif ( $codon =~ /AA[TC]/i) { return 'N' }
	elsif ( $codon =~ /CC./i) { return 'P' }
	elsif ( $codon =~ /CA[AG]/i) { return 'Q' }
	elsif ( $codon =~ /CG.|AG[AG]/i) { return 'R' }
	elsif ( $codon =~ /TC.|AG[TC]/i) { return 'S' }
	elsif ( $codon =~ /AC./i) { return 'T' }
	elsif ( $codon =~ /GT./i) { return 'V' }
	elsif ( $codon =~ /TGG/i) { return 'W' }
	elsif ( $codon =~ /TA[TC]/i) { return 'Y' }
	elsif ( $codon =~ /TA[AG]|TGA/i) { return '*' }
	elsif ( $codon =~ /N/i ){ return 'X' }
	else {
		#print STDERR "Bad codon \"$codon\"!!\n";
	}
}
```
