## Supplementary material for "Unique evolutionary radiation of odorant receptors in birds": DataS15_step_5_a_Start_M_pick_up.120.pl

```
#!/usr/bin/perl -w


my $foldername='./';
############ YOU HAVE TO SET THIS VALUE and ENTER FILE NAME FOR EACH SPP SEPARATELY ###########
my $first_tm=48;
# elut: 268
# mfur: 298
# pbra: 
# you get this from your mafft alignment after you remove bad sequences and realign
# as many times as necessary until no more sequences have >5 gaps in a TM domain
# then in the alignment, record the AA location of the start of the first domain
# note that it'll differ from alignment to alignment because of how much gappy sequences
# come before your first domain.

opendir(FOLDER,"$foldername") or die print "can not find the file";
my @array = grep(/step_4_result_mafftAlignment_einsi_filtered.fasta$/,readdir(FOLDER));
close FOLDER;


foreach my $filename ( @array )
{

my $input=$filename;
my $output="step_5_result.pickedM.fasta";
   ### $output=~s/\.fasta$/_M_picked.fasta/;

open (FILE,"$input");

open (OUT, ">>$output");

my @array; 

     my $species;
     my %sequences;
	
        while( <FILE> )	{
		    my $line=$_;  chomp $line;
                if( $line =~ /^>(.+)/ ){
                        $species = $1;
                        $sequences{$species} = '';  push(@array, $species) ;
                }
                else{
                            $sequences{$species} .= $line;                      
                   }

	 }
	close FILE;
	
	foreach my $name( @array)
	{ 
	    my $lengthall=length($sequences{$name});
         
		my $part2=substr($sequences{$name}, ($first_tm-1), ($lengthall-$first_tm+1));
           $part2=~s/-//g;		
	
	    my $Nsite=substr($sequences{$name}, 0, ($first_tm-1));
		$Nsite=~s/-//g;
		$Nsite=~s/\n//g;
		
		if ($Nsite){
		
		my $length=length($Nsite);
		my @array35; my @array2034; my @array21;
		my $loc=0;
				
		for (my $i=($length-1); $i>=0; $i--){
		$loc++;
		my $aa=substr($Nsite, $i, 1); # print OUT $aa,"\n";
		if ($aa=~/M/){
		if (($loc>20) and ($loc<35)) {push (@array2034,$loc)}
		if ($loc>=35) {push (@array35, $loc)}
		if ($loc<=20) {push (@array21, $loc)}
		            }	
        					
		}
		
		my $best_start;
		
		#print OUT @array2034,"\n";
		#print OUT @array35,"\n";
		#print OUT @array21,"\n";
		print OUT ">$name\n";		
		#print OUT $Nsite,"------------------\n";
		#print OUT $part2,"\n";
				
		if (@array2034)  {
		$best_start=min(@array2034);
		my $Nsequence=substr($Nsite, ($length-$best_start), $best_start);
		print OUT $Nsequence;
		print OUT $part2,"\n\n";
		#print OUT $best_start, "===============================================\n\n\n";
		next;}
		
		elsif (@array35)  {
		$best_start=min(@array35);
		my $Nsequence=substr($Nsite, ($length-$best_start), $best_start);
		print OUT $Nsequence;
		print OUT $part2,"\n\n";
		#print OUT $best_start, "===============================================\n\n\n"; 
		next;}
		
		elsif (@array21)  {
		$best_start=max(@array21);
		my $Nsequence=substr($Nsite, ($length-$best_start), $best_start);
		print OUT $Nsequence;
		print OUT $part2,"\n\n";
		#print OUT $best_start, "===============================================\n\n\n"; 
		next;}
		
	 }
        
	}
}
	
	
	
		  
	  sub max {
   my($max_so_far) = shift @_;
   foreach (@_){                         
      if($_>$max_so_far){                  
           $max_so_far=$_;
      }
   }
   return $max_so_far;                      
}

   

sub min {
   my($min_so_far) = shift @_;
   foreach (@_){                         
      if($_<$min_so_far){                  
           $min_so_far=$_;
      }
   }
   return $min_so_far;                      
}
		
```
