## Supplementary material for "Unique evolutionary radiation of odorant receptors in birds": DataS20_Filter_ORs.R

```
#### My version of Amber's olfaction cleaning script
require(GenomicRanges)
setwd("/media/balalab/SmallArseRAIDbac/ORs/genomes/All/Xenopus/RhiDar")
#setwd("/media/balalab/NewSmallRAID/lovebird/LB_RNAseq_for_Robert/RhiDar")

spp="RhiDar"
########### read back in the deduped (DD) results that you just wrote out (once) #########
blastsumDD_raw <- read.table("OR_query_mini_ORN_gharial.sum",header=T,sep="/",quote="")

blastsumDD_raw <- read.table("OR_query_mini_ORN_gharial.TAIL.sum",header=T,sep="/",quote="")

#blastsumDD_raw <- read.table("OR_query_mini_ORN_gharial.sum",header=T,sep="/",quote="")


# do some filtering ahead of time to speed up processing:
# get rids of hits < 250bp in length
tail(blastsumDD_raw)
dim(blastsumDD_raw) # 411298 
##### NOTE!!!! These lengths are in terms of Amino Acids but then when you do end - start it is nucleotides. So most should be > 810 NUCLEOTIDES (not AAs)
blastsumDD_raw$Hit_length <- as.numeric(as.character(blastsumDD_raw$Hit_length))
#blastsumDD <- blastsumDD_raw # - SPECIFICALLY FOR PSEUDO
blastsumDD <- blastsumDD_raw[blastsumDD_raw$Hit_length >= 250,] #- OMMITTED FOR PSUEDO
blastsumDD <- na.omit(blastsumDD)
blastsumDD$Hit_Start <- as.numeric(as.character(blastsumDD$Hit_Start))
blastsumDD$Hit_END <- as.numeric(as.character(blastsumDD$Hit_END))
#Hit_strand <- as.numeric(blastsumDD$Hit_strand)

dim(blastsumDD) # 318573

# want to add a unique index to each hit so that I can grab things out later.
blastsumDD$uniqueHitLabel <- seq(1:dim(blastsumDD)[1])

# 1. need to remove overlapping hits (within 100bp of each other) by choosing the one with the best e-value (lowest) or bit score (highest)
head(blastsumDD) 
# so what matters is Hit (scaffold ID), Hit_Start, Hit_End, Evalue and Bit Score of each hit.
#source("https://bioconductor.org/biocLite.R")
#biocLite("GenomicRanges")

# seqnames are chromosomes/scaffolds
# ranges use IRanges to give starts then ends 
# strand you get from results
# score is bit score (or could be evalue)
#write.csv(as.data.frame(blastsumDD), file="blastsumDD_test.csv")

grobj <- GRanges(seqnames=blastsumDD$Hit,ranges=IRanges(start=blastsumDD$Hit_Start,end=blastsumDD$Hit_END),strand=blastsumDD$Hit_strand,bit.score=blastsumDD$Bit.score,evalue=blastsumDD$E.value,query=blastsumDD$Query,hit.length=blastsumDD$Hit_length,uniqueHitLabel=blastsumDD$uniqueHitLabel)
# for some reason, need that space between Query 1; Mamu whatever
head(grobj)
########## Find overlaps, maxgap=100,minoverlap=1,drop hits to itself, drop redundant hits #######
# overlaps <- findOverlaps(grobj,maxgap=100L,minoverlap=1L,drop.self=F,drop.redundant=F)
overlaps <- findOverlaps(grobj,maxgap=100L,minoverlap=0L,drop.self=F,drop.redundant=F) # don't need redundant hits 

length(overlaps) # withonly length filter: 110610271

queryNums <- unique(queryHits(overlaps))

# "can use maxgap "intervals with a separation of maxgap or less and a minimum of minoverlap overlapping positions, allowing for maxgap, are considered to be overlapping. maxgap should be a scalar, non-negative, integer. minoverlap should be a scalar, positive integer." (Iranges manual)

# "In this case, and only this case, the drop.self and drop.redundant arguments are allowed. By default, the result will contain hits for each range against itself, and if there is a hit from A to B, there is also a hit for B to A. If drop.self is TRUE, all self matches are dropped. If drop.redundant is TRUE, only one of A->B and B->A is returned." (Iranges manual)
# start with index 1 (group by index)
# for q in queryhits ...
# put in your genomic ranges object, and your overlaps result:

############################################ RUN PRUNING FUNCTION #######################
cleanByEValue_prune <- function(grobj,overlaps) {
  queryNums <- unique(queryHits(overlaps))
  keep <- GRanges()
  while(length(queryNums) > 0){
    q <- queryNums[1]
    print(q)
    subs_q <- subjectHits(overlaps[queryHits(overlaps)==q]) # don't need to add the query number in because I have the hits to itself kept into overlaps now     
    grobj_subs <- grobj[subs_q]
    keepThisHit <- which.min(grobj_subs$evalue)
    keep <- c(keep,grobj_subs[keepThisHit])
    queryNums <- queryNums[!(queryNums %in% subs_q)]
    print(paste("dropped:",subs_q))
    print(paste("left to go: ", length(queryNums)))
  }
  return(unique(keep))
}

keep_prune_eval <- cleanByEValue_prune(grobj,overlaps)

# check it! 
check <- findOverlaps(keep_prune_eval,maxgap=100L,minoverlap=0L,drop.self=F,drop.redundant=F)
# should only be self:self matches left! 

# write out
pullOut <- blastsumDD[blastsumDD$uniqueHitLabel %in% keep_prune_eval$uniqueHitLabel,] # pull out the original entries 
############################# WRITE OUT .sum and .bed #####################################
write.table(pullOut,"cleanedByEvalue.LengthFilterOnly.Pruned.RhiDar_control.TAIL.sum",row.names = F,quote=F,sep="/")
pullOut$bed_HitStart0 <- pullOut$Hit_Start - 1 # because bed is zero based (don't change end bc it is non inclusive)
# want to add 300 (100 AAs) to the start and end (doesn't matter what strand because adding in both directions)
pullOut$bed_HitStart0_minus300 <- pullOut$bed_HitStart0 - 300 # if goes negative, set to 0
pullOut$bed_HitStart0_minus300[pullOut$bed_HitStart0_minus300 < 0] <- 0
pullOut$bed_HitEnd_plus300 <- pullOut$Hit_END + 300 # some may be over limit of chromosome -- deal with that when it happens 
#### check to make sure you don't overhang:

#In Unix run:
# samtools faidx Mustela_putorius_furo.MusPutFur1.0.dna.toplevel.fa &

#pullOut <- read.table("GCF_017654675.1_Xenopus_laevis_v10.1_genomic.fna.fai",header=T,sep="/",quote="")
# need to do samtools faidx on genome:
scaffLengths <- read.table("GCF_050947455.1_aRhiDar2.hap1_genomic.fna.fai",header=F)
colnames(scaffLengths) <- c("Scaffold","length")
head(scaffLengths)
pullOut_lengths <- merge(pullOut,scaffLengths[,c("Scaffold","length")],by.x="Hit",by.y="Scaffold",all.x=T,all.y=F)
head(pullOut_lengths)
# check for overhangs:
pullOut_lengths[pullOut_lengths$bed_HitEnd_plus300 > pullOut_lengths$length,]
# these are the bad ones that need to be fixed
pullOut_lengths[pullOut_lengths$bed_HitEnd_plus300 > pullOut_lengths$length,]$bed_HitEnd_plus300 <- pullOut_lengths[pullOut_lengths$bed_HitEnd_plus300 > pullOut_lengths$length,]$Hit_END

# check for overhangs again:
pullOut_lengths[pullOut_lengths$bed_HitEnd_plus300 > pullOut_lengths$length,]
#none! good! 
##### continue on:
# get strand:
pullOut <- pullOut_lengths
pullOut$bedstrand[pullOut$Hit_strand=="-1"] <- "-"
pullOut$bedstrand[pullOut$Hit_strand=="1"] <- "+"
# for the bedtools name want to have it be like Gang Li's script:
# $species\_$chr\_$start\_$end\_$string\ *** NOTE THAT BLAST IS ONE BASED *** so start end are 1 based
pullOut$bedname <- paste(spp,pullOut$Hit,pullOut$Hit_Start,pullOut$Hit_END,pullOut$Hit_strand,sep="_")
bed <- pullOut[,c("Hit","bed_HitStart0_minus300","bed_HitEnd_plus300","bedname","E.value","bedstrand")]
bed
dim(bed) # 716
write.table(bed,"RhiDar_control_ORs_OG.cleanedByEvalue.LengthFilterOnly.Pruned.scaff.0based.start.minus300.stop.plus300.name.eval.strand.TAIL.bed",row.names = F,col.names = F,quote=F,sep="\t")
```
